# Structure-guided engineering of α-ketoisocaproate dioxygenase increases isobutene production in *Synechocystis* sp. PCC 6803

**DOI:** 10.1101/2024.12.20.629387

**Authors:** Conrad Schumann, Amit Kugler, Bhavik Shah, Gustav Berggren, Henrik Land, Cecilia Blikstad, Karin Stensjö

**Author notes:** These authors contributed equally (Conrad Schumann, Amit Kugler).

## Abstract

Isobutene is a promising precursor for jet fuel due to its high energy density and favorable combustion properties. Light-driven bioproduction of isobutene has recently been investigated as an alternative strategy to crude oil refinement or fermentation-based manufacturing processes by harnessing the unicellular cyanobacterium *Synechocystis* sp. PCC 6803 and the α-ketoisocaproate dioxygenase (*Rn*KICD) from *Rattus norvegicus*. However, the obtained production level was not sufficient, partially due the promiscuous activity of *Rn*KICD. The enzyme catalyzes both the reaction with ρ-hydroxyphenylpyruvate (HPP) for homogentisate formation, as well as the reaction with α-ketoisocaproate (KIC), the precursor for isobutene synthesis. Here, to overcome this bottleneck step in the isobutene biosynthesis, protein engineering was employed to improve *Rn*KICD activity and in vivo isobutene production. Purified *Rn*KICD variants were characterized by measuring in vitro KIC and HPP consumption rates, as well as isobutene formation rate. The active site mutations F336V, N363A altered the KIC and HPP consumption rates, while the KIC-to-isobutene conversion ratio was only marginally affected. Besides, the *Rn*KICD variants F336V, N363A and F336V/N363A exhibited a substantially enhanced substrate selectivity for KIC over HPP. Among the examined engineered *Synechocystis* strains, Syn-F336V showed a 4-fold improvement in isobutene production, compared to the base strain (Syn-*Rn*KICD). Our findings reveal that residues F336 and N363 play a crucial role in substrate interactions, as targeted mutations at these sites shifted the substrate selectivity towards KIC and elevated the in vivo isobutene production levels significantly. We conclude that engineering the active site of *Rn*KICD is a potent tool for improving isobutene bioproduction in *Synechocystis*.

**Highlights:**

1. Promiscuous α-ketoisocaproate dioxygenase (KICD) from *Rattus norvegicus* was overproduced in *Synechocystis* sp. PCC 6803.
2. Protein engineering was used to improve substrate selectivity.
3. The generated F336V variant led to a 4-fold increase in isobutene production *in vivo*.

## Introduction

Isobutene is a volatile C4-alkene, serving as platform molecule for synthesizing numerous industrial products, such as butyl rubber, terephthalic acid, specialty chemicals and a gasoline performance additive (Marchiona et al., 2001). In addition, isobutene can be used as precursor for the manufacturing of jet-fuels, for which isobutene undergoes oligomerization and hydrogenation reactions (Nicholas, 2017). At present, isobutene is primarily produced by extraction from steam cracking of petroleum hydrocarbons (Gholami et al., 2021).

Biofuel manufacturing by microorganisms offers a promising approach to support a circular bioeconomy. The environmental impact of a production can be reduced by recovering the remaining value of by-products or waste, e.g. using captured CO_2_ in the production of valuable compounds like isobutene (Tcvetkov et al., 2019). As a volatile gas, isobutene can be efficiently collected and purified from off-gas, minimizing downstream processing costs and risk of in vivo accumulation of the fuel compound to a toxic level.

We have recently described the light-driven biosynthesis of isobutene from CO_2_ in the model cyanobacterium *Synechocystis* sp. PCC 6803 (hereafter *Synechocystis*) (Mustila et al., 2021). This was achieved by metabolic engineering through heterologous expression of the gene encoding α-ketoisocaproate dioxygenase from *Rattus norvegicus* (*Rn*KICD; EC 1.13.11.27) However, the low isobutene yields make this system economically less feasible compared to fermentative processes that use sugars as feedstocks (Gogerty & Bobik 2010., van Leeuwen et al., 2012) and limits its application/operation at an industrial scale (Mustila et al., 2021).

Several reported enzymes involved in biocatalytic production of isobutene catalyze the irreversible decarboxylation of a C5 branched chain carboxylic acid (Fukuda et al., 1994, Gogerty & Bobik, 2010, Rossini et al., 2015). The catalyzed reaction is either facilitated via oxidative, ATP-dependent (Gogerty & Bobik, 2010, Rossoni et al., 2015) or redox active cofactor dependent decarboxylation (Fukuda et al., 1994, Saaret et al., 2021). The mononuclear Fe(II)-dependent *Rn*KICD has been reported to catalyze the conversion of α-ketoisocaproate (KIC) and molecular oxygen into 3-hydroxy-3-methylbutyrate (HMB) (Sabourine & Bieber, 1982, 1983) followed by a non-enzymatic decomposition reaction to yield isobutene (Rossini et al. 2015). In our previous study, we found that in vitro isobutene production rate exceeded the isobutene formation rate from spontaneous HMB decomposition. This demonstrates that isobutene can be directly converted from the C6 branched chain carboxylic acid KIC by undergoing two decarboxylation steps (Mustila et al., 2021). Hence, *Rn*KICDs promiscuous nature of converting KIC results either in the formation of HMB or isobutene, with low product specificity for isobutene. However, the mechanism underlying the isobutene formation remains unclear.

*Rn*KICD shares a 100% amino acid sequence identity with the rat ρ-hydroxyphenylpyruvate dioxygenase (HPPD) which converts ρ-hydroxyphenylpyruvate (HPP) into homogentisate (HGA) (Baldwin 1995, Sabourine & Bieber, 1982, 1983). Thus, mammalian HPPD and KICD is the same enzyme that displays substrate promiscuity towards a variety of α-keto acids (Crouch et al., 1997). In plants, HGA acts as a precursor for plastoquinone-9 and α-tocopherol synthesis, which are important for the photosynthesis process and plant survival (Norris et al., 1995). Due to the essential physiological function of HPPD, this enzyme has been intensively investigated as a target for herbicide discovery (Wang et al., 2015).

Protein engineering is an important part of metabolic engineering which involves amino acid substitutions for developing enzymes with increased catalytic rate, substrate specificity or stability (Bilal et al., 2018). In turn, such modifications can enhance titers and yields of the compound of interest (Jamil et al., 2022). Protein engineering strategies include directed evolution (random mutagenesis), rational design (site-directed mutagenesis), or a combination of both (semi-rational) (Xiong et al., 2021). The choice of approach depends on existing knowledge of the enzyme being studied and the availability of screening or selection methods for the variants produced. Recent crystallographic data has identified key residues within the HPPD binding pocket, involved in substrate (HPP) binding and catalytic efficiency (Huang et al., 2021), which can assist in repurposing HPPD activity by rational and semi-rational designs.

In this study, we generated *Rn*KICD variants with active site amino acid substitutions based on rational and semi-rational protein design. This with the aim to increase selectivity towards KIC as isobutene precursor, and ultimately enhance the isobutene production from cultivated *Synechocystis*. All engineered *Rn*KICD enzyme variants displayed lowered in vitro consumption rates for the two substrates HPP and KIC, while the apparent selectivity towards KIC was increased. The generated strain Syn-F336V overexpressing *Rn*KICD-F336V showed a 4-fold higher isobutene titer, compared with the base strain. Our results demonstrate that protein engineering of *Rn*KICD is a feasible strategy to significantly increase the production of isobutene synthesized by metabolically-engineered *Synechocystis*.

## Results

### Substrate promiscuity of RnKICD limits isobutene synthesis

We previously described the recombinant production of α-ketoisocaproate dioxygenase (*Rn*KICD) in *Synechocystis* sp. PCC 6803 for the conversion of KIC into isobutene (Mustila et al., 2021). However, the isobutene synthesis is limited due to the promiscuous activity of *Rn*KICD, accepting both KIC and HPP as substrates (Figure 1). Since HPP can act as a competetive inhibitor for the KIC conversion, we hypothesized that this substrate promiscuity creates a bottleneck for the isobutene in vivo biosynthesis. In this work, we applied rational and semi-rational protein engineering on residues in the active site coordination sphere to improve substrate selectivity and thereby in vivo isobutene production.

**Figure 1.**
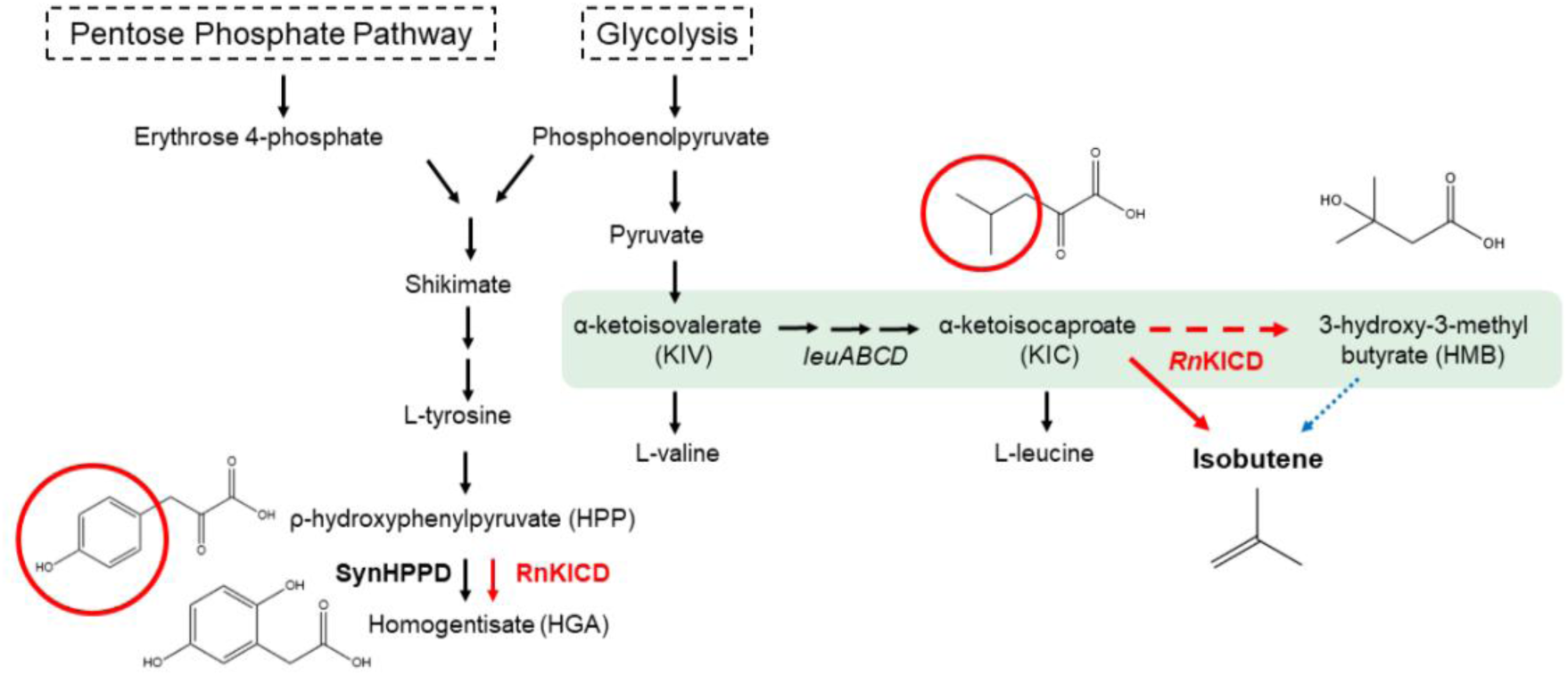
Involvement of *Rn*KICD in the endogenous homogentisate and heterologous isobutene biosynthesis pathways. *Rn*KICD utilizes α-ketoisocaproate (KIC) to produce isobutene, and ρ-hydroxyphenylpyruvate (HPP) to produce homogentisate. Black arrows indicate native enzymes; solid and dashed red arrows indicate catalysed non-native reactions; blue dotted arrow inicates non-ezymatic decomposition. Red circles indicate structural differences between KIC and HPP.

### Selection of amino acid residues in RnKICD for protein engineering

The limited mechanistic knowledge of *Rn*KICD catalyzing the conversion of KIC represents a challenge for a rational design approach. Yet, the binding of HPP, as well as the catalytic mechanisms of HPPDs (*Rn*KICD among them), have been elucidated in several studies (Huang et al., 2021, Lin et al., 2019). As reported, the main active site residues of HPPD involved in HPP binding are Q251, Q265, Q334, F336 and N363 (Figure 2 A). In the open conformation, the phenolic hydroxyl of HPP forms a hydrogen bond with the side-chain of N363, which helps to direct the substrate into the active site. Then, the substrate flips and the phenol ring instead partakes in hydrogen bonding with Q251 and Q265 as the active site closes. The benzene ring of HPP is also involved in T-π stacking (edge-to-face) with the phenyl sidechain of F336. Another glutamine residue, Q334, forms a hydrogen bond with the pyruvate moeity of HPP which also interacts with the active site ferrous ion.

**Figure 2.**
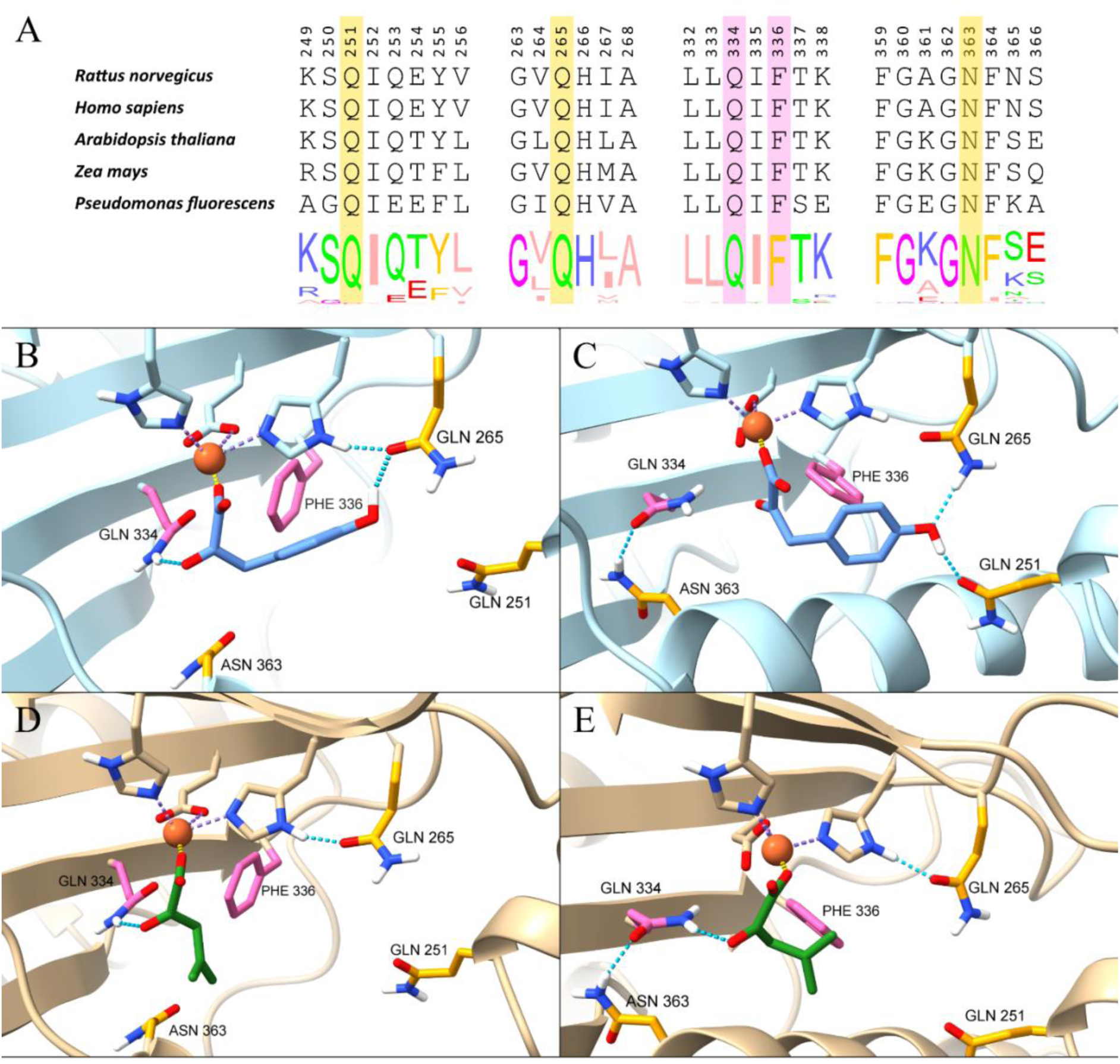
Sequence and structural analysis of HPPD enzymes. A. Alignment of *Rn*KICD amino acid sequence with selected sequences of HPPD enzymes from various species and consensus logos showing the conservation within each position. Sections not relevant to this study are omitted for clarity. Normalized consensus logos were generated in Jalview based on the 262 sequences used for multiple sequence alignment. B. Homology model of the *Rn*KICD active site in open conformation with HPP as substrate. C. Homology model of the *Rn*KICD active site in closed conformation with HPP as substrate. D. Homology model of the *Rn*KICD active site in open conformation with KIC as substrate. E. Homology model of the *Rn*KICD active site in closed conformation with KIC as substrate. Rational design targets are highlighted in yellow, and the site-saturation targets in pink. The structure of HPP is shown in light blue, and the structure of KIC in green. The iron atom is represented by an orange sphere.

To analyze the active site residues of *Rn*KICD and its interactions with the substrates HPP and KIC, docking simulations were performed with homology models of *Rn*KICD using both open and closed conformations (Figure 2 B-E). In docking studies of HPP with homology models of *Rn*KICD, similar binding/interactions of HPP with the active site residues were observed as reported in the literature (Figure 2 B-C). In the open conformation, two main binding modes of HPP were observed: 37/50 dockings resulted in hydrogen bonding between HPP and Q265 (Figure 2 B) and 13/50 dockings resulted in hydrogen bonding between HPP and N363 (Supplementary Figure 1). Regarding KIC as substrate, we found that KIC is mainly involved in electrostatic interaction with the ferrous ion and hydrogen bonding with Q334 and a probable hydrophobic interaction with F336 (Figure 2 D-E).

We targeted F336 for mutagenesis with the hypothesis that substitution of F336 would disrupt the T-π stacking between HPP and F336 and thus reduce the rate of HPP conversion. Furthermore, we targeted Q334 with the aim of reducing formation of the enzyme-HPP complex. Since KIC also interacts with Q334 (Figure 2 C and D), it is plausible that the introduction of any mutation at the Q334 position would also effect the interaction with KIC. To find an optimum mutation for F336 and Q334 we used site directed saturation mutagenesis. In addition, we used rational design to examine the effect of the mutations Q251E, Q265E and N363A within the active site. We hypothesized that the replacement of glutamine with glutamate (Q251E and Q265E) would introduce an electrostatic repulsion effect towards HPP, while retaining a similar structural composition of the active site. The substitution of asparagine with an alanine at position 363 would decrease HPP binding due to disruption of hydrogen bonding and might also result in favored KIC binding, due to an increased hydrophobic interaction.

### Screening of site-saturation mutant libraries

To evaluate the expected nucleobase distribution of the site-saturation libraries, a quick quality control (QQC) sequensing protocol was used, targeting the codons for Q334 and F336 (Kille *et al.,* 2013). The QQC matched in both cases qualitatively the expected nucleobase distribution with exclusively T or G in the third codon position, while including all four nucleobases in the first and second position (Supplementary Figure 2). Recombinantly-produced *Rn*KICD variants were thereafter screened for altered substrate selectivity compared to wild type (WT) enzyme. Hits from both libraries were selected based on (I) high KIC consumption activity and (II) altered substrate consumption for one or both substrates (Figure 3). Additionally, sequencing was performed and translated amino acids in each variant identified (Supplementary Figure 3).

**Figure 3.**
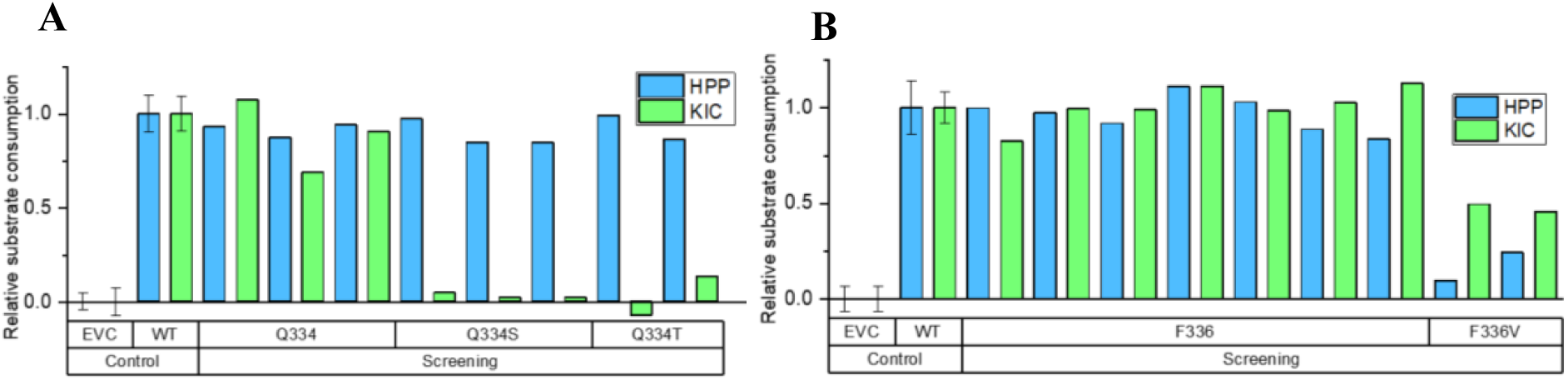
Selected hits from the activity screens of transformant libraries generated by site-saturation mutagenesis. Potential hits for the F336 and Q334 libraries were selected based on two individual criteria: (I) high KIC consumption and (II) altered consumption for one or both substrates. A. Screening result for the selected colonies from the Q334X mutant library. Relative consumption rate of KIC (green) and HPP (blue) by Q334X mutants in comparison to WT with the introduced mutation below. B. Screening result for the selected colonies from the F336X mutant library. Relative consumption rate of HPP (blue) and KIC (light green) by F336X mutants in comparison to WT with the introduced mutation below. WT and empty vector control (EVC) for comparison are shown as average with a standard deviation from eight wells per 96-well plate.

All selected variants with wildtype-like high KIC consumption (criterion I) were found to be expressing non-altered *Rn*KICD variants, highlighting the importance of Q334 and F336 for high consumption of KIC. Two F336 library mutants displayed altered substrate consumption (criterion II), with mutation F336V enhancing selectivity towards KIC. In this mutant, HPP consumption was reduced about 7-fold compared to WT, while KIC consumption decreased only 2-fold (Figure 3B). The slight reduction in KIC consumption was considered a side effect of the mutation. From the Q334 mutant library, five mutants were selected based on criterion II, showing WT *Rn*KICD-like HPP consumption but disrupted KIC consumption. These results suggest that mutations Q334S and Q334T contributed to increased HPP selectivity, reinforcing the importance of Q334 in KIC consumption by *Rn*KICD (Figure 3A).

### In vitro analysis of RnKICD and generated variants

The previously mentioned rationally designed variants (Q251E, Q265E and N363A) and the best performing variant from the semi-rational design (F336V) were purified (Supplementary Figure 6) and assessed with regards to consumption of substrates KIC and HPP, as well as formation of isobutene (Supplementary Figure 4). We found that Q251E and Q265E resulted in practically sluggish *Rn*KICD variants. Both mutations decreased HPP consumption by approximately 10-fold while the KIC consumption was too small to measure. (Figure 4A). Consequently, the in vitro measurements for these variants resulted in no detectable isobutene formation (data not shown). In contrast, N363A had comparable KIC consumption rate and isobutene formation to that of WT *Rn*KICD, while the HPP consumption was ∼10-fold lower (Figure 4 A & B). Hence, the selectivity towards KIC was enhanced by the N363A amino acid substitution. For F336V, the consumption rate of HPP was critically impaired, while the KIC consumption and isobutene production rates were 35% and 40% lower as compared to WT, respectively (Figure 4 A-B).

**Figure 4.**
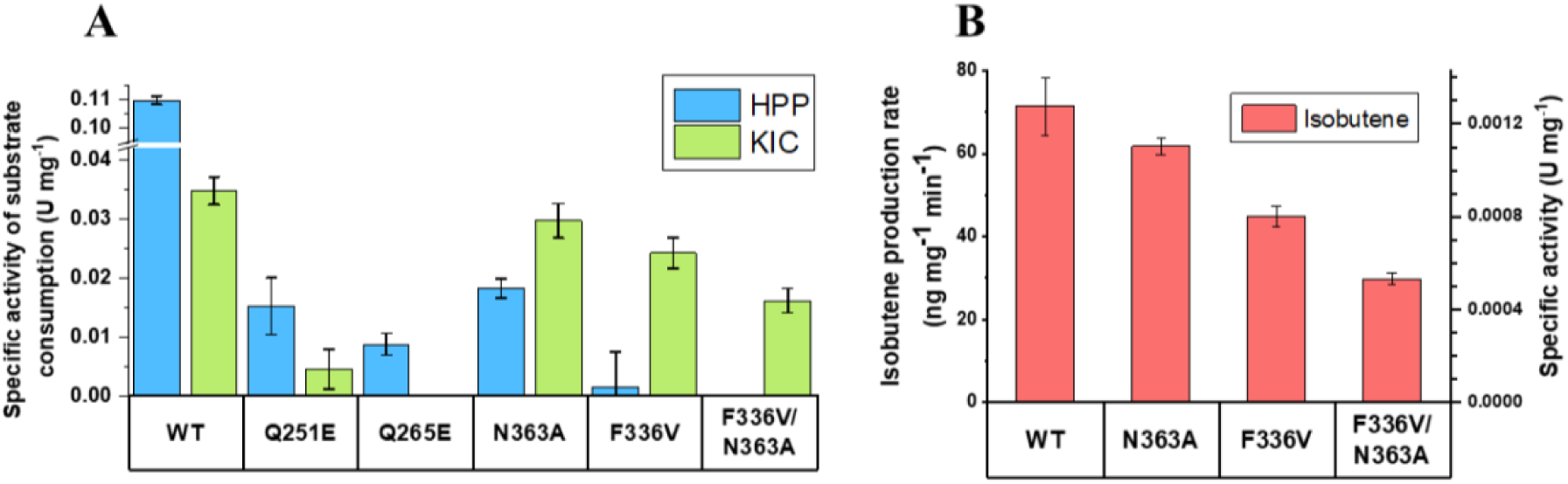
Functional characterization of *Rn*KICD variants. A. KIC and HPP consumption rate at 2.5 mM substrate expressed as specific activity (U mg^-1^). B. Isobutene production rate (ng mg^-1^ min^-1^) at 3 mM KIC also expressed as specific activity (U mg^-1^). U = µmol min^-1^.

The two single-point mutations that retained KIC consumption activity (F336V, N363A) resulted in different characteristics compared to WT *Rn*KICD. N363A exhibited an improved selectivity towards KIC with retained isobutene production, while F336V showed even higher selectivity, but with decreased KIC consumption and isobutene production. In order to explore any synergistic effects between the two mutations, the double mutant F336V/N363A was created. This new variant displayed a complete disruption of the HPP consumption whereas the KIC consumption and isobutene production were both retained at lower levels, as compared to the single-point mutations. Yet, comparing the specific activities of isobutene formation and KIC consumption revealed that all three variants exhibited similar KIC-to-isobutene conversion ratios (3.3-3.7%), comparable to the wild type (Supplementary Table 3).

### In vivo analysis of engineered Synechocystis strains

The results obtained by the in vitro assays revealed a higher substrate specificity for some of the *Rn*KICD variants, leading to a potential for improving isobutene production in vivo. Therefore, the genes encoding WT *Rn*KICD and the best performing mutant variants were individually introduced into *Synechocystis*, resulting in the strains Syn-KICD, Syn-N363A, Syn-F336V and Syn-F336V/N363A, respectively (Table 1).The performance of *Synechocystis* strains producing the different *Rn*KICD variants was evaluated in terms of growth and final isobutene titer (Figure 5). No significant growth differences were observed among the examined *Synechocystis* strains (Figure 5 A), indicating that the production of *Rn*KICD did not pose a metabolic burden on the cell. After four days of batch cultivation in GC-MS vials (Figure 5 B), strains Syn-KICD and Syn-F336V/N363A showed a comparable isobutene production, while the isobutene production by Syn-N363A was decreased significantly. However, Syn-F336V exhibited a considerable increase, comprising a 4-fold enhancement of absolute isobutene titres as well as growth-related titeres, as compared to the base strain (Syn-KICD) (Figure 5 B and Supplementary Figure 5).

**Figure 5.**
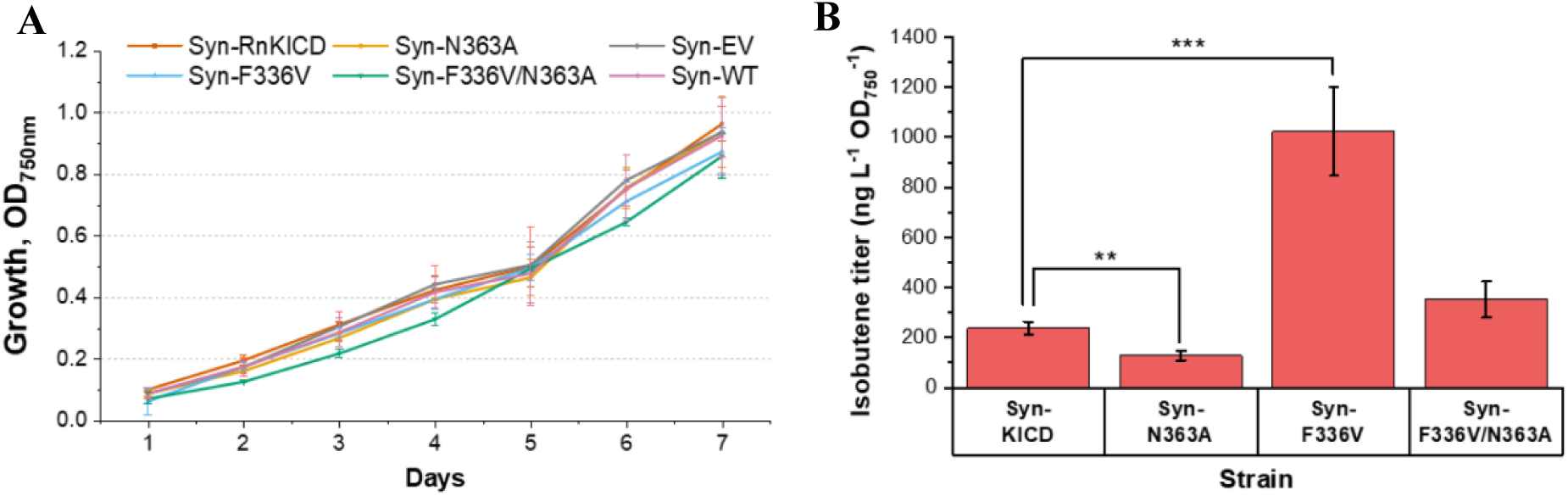
Production capabilites of *Synechocystis* strains overexpressing *Rn*KICD and its variants after four days of batch cultivation. A. Growth of isobutene-producing strains, as indicated by optical density (OD_750nm_). B. Isobutene production titer (ng L^-1^ OD_750_^-1^) normalized to the culture density (OD_750nm_). All the results represent the mean of five biological replicates; error bars represent the standard deviation. Asterisks represent significant differences between the corresponding strain and the base strain, ** p < 0.01, *** p < 0.001 in *t*-test.

**Table 1.** Strains and genetic constructs used in this study.

| Strain | Construct | Description | Reference |
| --- | --- | --- | --- |
| Syn-EVC | pEEK2-Km<sup>R</sup> | *Synechocystis* sp. PCC 6803 strain harboring pEEK2 plasmid for Km resistance as empty vector control | Mustila et al., 2021 |
| Syn-RnKICD | pEEK2-RnKICD | *Synechocystis* sp. PCC 6803 strain harboring pEEK2 plasmid for Km resistance and RnKICD | Mustila et al., 2021 |
| Syn-N363A | pEEK2-N363A | *Synechocystis* sp. PCC 6803 strain harboring pEEK2 plasmid for Km resistance and N363 mutation in RnKICD | This study |
| Syn-F336V | pEEK2-F336V | *Synechocystis* sp. PCC 6803 strain harboring pEEK2 plasmid for Km resistance and F336V mutation in RnKICD | This study |
| Syn-N363A/F336V | pEEK2-N363A/F336V | *Synechocystis* sp. PCC 6803 strain harboring pEEK2 plasmid for Km resistance and N363/F336V mutation in *RnKICD* | This study |
| EcBL21-EVC | pETBB-Km<sup>R</sup> | *E. coli* BL21(DE3) strain harboring pETBB plasmid for Km resistance as empty vector control | Mustila et al., 2021 |
| EcBL21-RnKICD | pETBB-*RnKICD* | *E. coli* BL21(DE3) strain harboring pETBB plasmid for Km resistance and *RnKICD* | Mustila et al., 2021 |
| EcBL21-Q251E | pETBB- Q251E | *E. coli* BL21(DE3) strain harboring pETBB plasmid for Km resistance and Q251E mutation in *RnKICD* | This study |
| EcBL21- Q265E | pETBB- Q265E | *E. coli* BL21(DE3) strain harboring pETBB plasmid for Km resistance and Q265E mutation in *RnKICD* | This study |
| EcBL21-N363A | pETBB-N363A | *E. coli* BL21(DE3) strain harboring pETBB plasmid for Km resistance and N363A mutation in *RnKICD* | This study |
| EcBL21-F336V | pETBB-F336V | *E. coli* BL21(DE3) strain harboring pETBB plasmid for Km resistance and F336V mutation in *RnKICD* | This study |
| EcBL21-N363A/F336V | pETBB-N363A/F336V | *E. coli* BL21(DE3) strain harboring pETBB plasmid for Km resistance and N363A/F336V mutation in *RnKICD* | This study |

## Discussion

In this study, semi-rational protein engineering of *Rn*KICD has been employed to enhance substrate specificity towards α-ketoisocaproate (KIC) in vitro and isobutene production titres in vivo. We identified two *Rn*KICD variants F336V and F336V/N363A that exhibited an improvement in substrate selectivity, yielding higher isobutene production in *Synechocystis*.

As previously shown by crystallographic and spectroscopic studies (Huang et al., 2021), the active site of HPPD is enclosed by a C-terminal α-helix, exhibiting two distinct orientations, assuming to function as a gate that controls the binding of the substrate in the active site (Lin et al., 2013). This helix is either in a closed conformation, shielding the active site from the solvent, or in an open conformation that allows access of substrate to the binding pocket. The catalysis is initiated by a bidentate interaction of HPP with the ferrous ion in the active site, hydrogen bonds with a conserved glutamine residue (Q334) and an association with the molecular oxygen, resulting in an octahedral coordination geometry involving a facial triad (H183, H266 and E349). This is followed by a keto acid moiety decarboxylation, phenyl ring hydroxylation, and side-chain re-arrangement.

### Rational and semi-rational protein design for identifying amino acids substitutions linked to altered substrate selectivity of RnKICD

Earlier studies show that by replacing the glutamines Q251 and Q265 to a glutamate in the active site of HPPD, the ability to form hydrogen bonds with the hydroxyl group of HPP was lost, thus decreasing the stability of the enzyme-substrate complex (Raspail et al., 2011). Therefore, we initially reasoned that introducing Q251E and Q265E mutations would reduce competition with KIC binding. However, our results indicate that these mutations not only impacted the HPP consumption, but also diminished KIC consumption. This highlights the importance of these glutamines for the structural integrity of the active site in *Rn*KICD. For future studies, substituting Q265 with smaller polar amino acids, such as serine or threonine might be a strategy to decrease the binding affinity of HPP whilst preserving the hydrogen bond with iron ligating residue H266.

The residue N363 is thought to play a role in the rotation of the C-terminal helix hinge region (Huang et al., 2021), which helps to bind HPP to the active site in its open conformation (Supplementary Figure 1). Previous studies demonstrated that the mutation N363A decreased *k*_cat_/*K*_m_ for HPP by 5-fold compared to the WT enzyme (Huang et al., 2021). Under our in vitro assay conditions, N363A showed similarly a 6-fold decrease in HPP consumption, while the KIC consumption was not affected. This change likely results from the decreased stabilization of HPP in the open conformation of *Rn*KICD. However, in vivo studies revealed that the Syn-N363A strain produced about half the amount, compared to the Syn-KICD strain. These findings suggest that while the N363A substitution significantly reduced HPP consumption in vitro, HPP may still be able to interact with the active site due to alternative binding (Figure 2B). Thus, HPP could still act as competitive inhibitor and thereby impact the isobutene production in *Synechocystis*.

In our semi-rational design library screening of F336X and Q334X, we aimed to identify hits with altered selectively towards one of the two substrates. We got two hits with increased selectivity towards HPP, Q334S and Q334T (Fig 3B). The loss of KIC activity in these mutants could be explained by the docking simulations on the *Rn*KICD homology model, showing that Q334 is probably essential for the KIC turnover, since it is the only amino acid directly interacting with KIC as substrate in both the open and closed conformations (Figure 2D and 2E).

Regarding the F336X variants, the absence of T-π stacking against the phenyl sidechain of F336 resulted in reduced activity towards HPP, whereas the activity towards KIC was not compromised to the same extent. This indicated that F336V is a promising candidate for improved isobutene production in vivo. Earlier kinetic experiments on HPPD showed that F336 does not directly participate in the catalysis of HPPD, but has importance for stabilizing the HPP-enzyme complex (Raspail et al., 2011). The replacement of F336 with nonpolar amino acids, such as valine, could increase interaction of KIC. It might be that alanine is too small to sustain substrate stabilization, whereas larger apolar amino acids, such as valine, seem to be beneficial for the formation of the enzyme-substrate complex. As the obtained in vitro isobutene formation rates were low in comparison to KIC consumption rates, it can be concluded that *Rn*KICD is catalytically promiscuous and isobutene is only formed as by-product, as previously discussed (Mustila et al., 2021).

### Targeting RnKICD substrate selectivity yields optimized biosynthetic pathway for isobutene production

This study aimed to enhance *Rn*KICD selectivity for KIC as isobutene precursor using rational and semi-rational protein design. The optimzed enzymes were assessed in vitro for their substrate consumption profiles, and strains exhibing a preference for KIC over HPP, were tested in vivo to strategically redirect biosynthesis toward increased isobutene production in *Synechocystis*. The goal was succesfully achieved, as strains Syn-F336V and Syn-F336V/N363A demonstrated improved isobutene production, with Syn-F336V producing 4-fold higher isobutene titer compared to base strain.

In *Synechocystis Rn*KICD acts in two distinct metabolic pathways resulting in a competition of KIC and HPP over the active sites. Hence, a decreased affinity for HPP reduces also the competitive inhibition for the KIC-to-isobutene conversion. Besides, the catalyzed reaction could be dependent on the cellular environment, such as substrate concentrations and pH. The contrary isobutene production rates from F336V and WT in vitro and in vivo suggest that environmental factors unique to the cellular context have a major influence enzymatic activity.

The in vitro assays do not account for cellular factors such as substrate competition of KIC and HPP or the availability of reducing co-substrates that help to reduce the mononuclear iron in the active site after one catalytic cycle. *Rn*KICD is using both KIC and oxygen as substrates, thus the *Rn*KICD activity might be limited by oxygen and KIC concentration under in vitro and in vivo conditions, respectively. The oxygen-rich cellular environment of a cyanobacterium could enhance the catalysis of the enzyme and thus explain at least part of the differential rates of isobutene production detected in vitro and in vivo.

## Conclusions

Using a semi-rational design, we first created an *Rn*KICD mutant library that was screened for enzyme variants with improved KIC-to-HPP consumption ratio. These enzymes were purified and characterized for improved isobutene formation. Different *Rn*KICD variants exhibited increased preference towards KIC as substrate and isobutene as product. Q251 and Q265 were found to be essential in retaining activity of *Rn*KICD independent of substrate. The introduced mutations in position Q334 disrupted the activity towards KIC, suggesting a distinct role of Q334 for KIC to isobutene formation. Further, we suggest that F336 and N363 are important for modulating selectivity between the two substrates HPP and KIC. Mutant strains of *Synechocystis* carrying the F336V variant increased isobutene production in vivo. The enhanced production of isobutene in the F336V strain underlines the importance of taking the in vivo context in consideration in designing in vitro enzymatic assays as a a tool for enhanced metabolite production. We conclude that a strategy of rational design and site-specific saturation mutagenesis within the active site of *Rn*KICD is a potent strategy for increasing substrate specificity and thus redirecting metabolic fluxes for improved isobutene bioproduction in *Synechocystis*.

## Materials and methods

### Bacterial strains and growth conditions

*E. coli* TOP10 (Invitrogen) were used for cloning, conjugation and screening for site-directed mutants, as well as assessing the plasmid mutant library quality. *E. coli* BL21 (DE3) was used for the heterologous overexpression of the generated RnKICD mutants. *E. coli* TOP10 cells were cultivated in lysogeny broth (LB) medium (Sigma Aldrich) or on 1.5% agar (w/v) plates, at 37 °C. *E. coli* BL21 (DE3) cells were cultivated in Terrific Broth (TB) medium (Sigma Aldrich), at 37 °C. All medium were supplemented with (50 μg mL^−1^ kanamycin).

The glucose-tolerant *Synechocystis* sp. PCC 6803 sub-strain (Trautmann et al., 2012) was used throughout the study. *Synechocystis* cells were cultivated in BG11 (Stanier et al., 1971) medium, or on 1.5% agar (w/v) plates, under 15 µmol photons m^−2^ s^−1^ at 30 °C. All *Synechocystis* cultures were supplemented with 50 μg mL^−1^ kanamycin. All strains used in this study are listed in Table 1.

### Homology modeling and molecular docking

To obtain structural models of both the open and the closed conformation of *Rn*KICD homology modeling was used. More specifically, a homology model in the open conformation was made using *At*HPPD (PDB ID: 5XGK) as template and a homology model in the closed conformation was made using *Hs*HPPD (PDB ID: 3ISQ) as template using YASARA (Krieger & Vriend 2014) structure version 18.3.23 following previously described protocols (Land & Humble 2018). Any ligands were removed from the resulting models and energy minimization was performed inside a water-filled simulation cell (cubiod shape, 5 Å around all atoms) with periodic boundaries using the AMBER 14 force field (Case et al., 2023). The energy minimized models were then subjected to docking simulations as previously described (Land & Humble 2018) using both HPP and KIC as ligands (50 docking runs per model/substrate combination). The most prevalent docking conformation out of the 50 runs for each docking simulation was chosen as the representative for further analysis.

### Bioinformatic analysis

In order to obtain a representative collection of sequences for bioinformatic analysis, separate BLASTp searches were performed using template sequences from different branches of life: *Rattus norvegicus* (Genbank ID: NP_058929.1), *Arabidopsis thaliana* (Genbank ID: NP_172144.3) and *Pseudomonas fluorescens* (Genbank ID: 1CJX_A). 2839 sequences were combined and trimmed to 262 sequences by removing 90% redundancy using Jalview (version 2.11.3.3). A multiple sequence alignment was then performed using ClustalΩ. (Sievers et al., 2014).

### Plasmid construction

The plasmids used in this study are detailed in Table 1. For heterologous expression in *E. coli* BL21 (DE3) plasmids pETBB-Km^R^, pETBB-*Rn*KICD plasmids previously constructed were used (Mustila et al., 2021). The self-replicating plasmids pEEK2-Km^R^ and pEEK2 were used for expression in *Synechocystis*. pEEK2-*Rn*KICD was previously constructed and contains a strong constitutive P*trc*_core_ and a codon optimized *kicd* (Mustila et al., 2021). Selected mutations were introduced into the *Rn*KICD gene on pETBB-Km^R^, and pEEK2-Km^R^. All primers used for site directed mutagenesis are listed (Supplementary Table 1). The coding sequence for Strep-tagged WT *Rn*KICD is listed as well (Supplementary Material 1).

All primers were synthesized and supplied by IDT (USA). The site-directed mutagenesis was carried out by an overlapping PCR-based method (Zheng 2004). The entire plasmids were amplified using Phusion Hot Start II High–Fidelity DNA Polymerase (Thermo Fisher Scientific), with subsequent digestion with DpnI enzyme (Shenoy & Visweswariah, 2003). The PCR products was then used for conjugation of *Synechocystis*.

### Site-saturation mutagenesis

The pETBB-*RnKICD* plasmid (Mustila et al., 2021) was used as the initial template for site-saturation mutagenesis and for the generation of two plasmid libraries with mutations in the positions Q334 and F336. For each mutant library four 5’-phosphorylated primers were designed and ordered by IDT (USA) (Supplementary Table 2). Mutations were introduced by miss-matches in the 5’-end of the primers, with degenerate primers allowing for various codons in one reaction (Wells et al., 1985). Using a primer mix with two degenerate (NDT and VHG) codons and one defined codon (TGG) limited introduced codons to 22 covering all 20 proteinogenic amino acid and excluding stop codons (Kille *et al*. 2013) This approach reduced genetic diversity, thus lowering the screening effort required. For the F336 library, forward primers contained the mutational codons, while revers primers for the Q334 library comprised the mutational codons as a reverse complement. The NDT:VHG:TGG primers for both libraries were mixed at 12:9:1 ratio (Kille et al., 2013) and used as forward and reverse primers in the PCR for F336 and Q334 mutant library, respectively. The T_m_ for all eight primers ranged between 62-65 °C (NEB Tm calculator, 2022).

Site-saturation mutagenesis used pETBB_*Rn*KICD_NStrepII (6.4 kb) as the template in eight 20 µL inverse PCRs (Ochman et al., 1988). For details, see Supplementary Table 2. Due to the variety of primers an annealing temperature gradient was applied to counteract preferential annealing. PCR products were gel purified, then incubated for 15 min at 37 °C with DpnI fast digest (Thermo Fischer) to remove template DNA, cleaned with a PCR clean-up kit (Zymo Research) and ligated overnight with Quick Ligase (New England Biolabs at room temperature (RT)). The ligation mix was used directly to transform *E. coli* TOP10 and *E. coli* BL21 (DE3) cells. Approximately 500 clones from each mutant library were pooled sequenced to evaluate the library quality based on introduced codons and the nucleobase distribution. This quick quality control (QQC) of the libraries confirmed effective template DNA removal and intended codon diversity, with mutations validate by Sanger sequencing (Mix2Seq, Eurofins Genomics).

### Protein production and purification

All *E. coli* culturing was done in terrific broth (TB) medium with kanamycin (50 μg mL^-1^). *E. coli* BL21 (DE3) cells (Invitrogen), harboring pETBB-RnKICD plasmids were cultured overnight in 20 mL in 100 mL Erlenmeyer flasks, at 37 °C and 190 rpm. The culture was then diluted to OD_600nm_ 0.1 and incubated in 1 L in a 3 L Erlenmeyer flask, at 37 °C and 190 rpm. Upon reaching OD_600nm_ 0.6-0.8 gene expression was induced with 0.5 mM, the temperature lowered to 20 °C, and incubation continued for 20 hours. Cells were harvested by centrifugation (Sorvall RC-3B Plus, GMI) at 4°C, 7,277 x g for 10 min and stored in −80 °C. Cells were lysed by resuspending in 50 mM HEPES (pH 7.5) with 1.2 mg mL^-1^ lysozyme, 0.06 mg mL^-1^ each of DNAse, RNAse, MgCl_2_ (2.4 mg mL^-1^), and EDTA free cOmplete™ protease inhibitor (Roche). After 30 min incubation, the suspension was sonicated on ice (Sonics Vibra Cell, CV33 tip) for 1 minute at amplitude 50% (10 seconds on and 20 seconds off). Soluble proteins were separated by ultracentrifuge (Culter Optima L-90K, rotor 70Ti, Beckman) (40 min, 4 °C, 257,000 x g, and filtrated through a 0.22 µm filter (Sigma-Aldrich).

Proteins were purified at 4°C using an ÄKTA pure FPLC system with two tandem Strep-Tactin Sepharose columns (StrepTrap HP 5 mL, Cytiva), with minor modifications to the manufacturer’s protocol using 50 mM HEPES (pH 7.5) as binding and 2.5 mM D-desthiobiotin in 50 mM HEPES (pH 7.5) as elution buffer. Purified proteins were concentrated with 30 kDa cut-off filters (Amicon, Merck Millipore) at 4 °C, 4,500 x g for 40 min. Protein purity was assessed by SDS-PAGE, and concentration was determined at 280 nm using a NanoDrop (Thermo Fisher) with and Molar extinction coefficient of 55,600 M cm^-1^ (ProtParam tool on the ExPASy Server, Gasteiger et al., 2005).

### Conjugation of Synechocystis sp. PCC 6803

*Synechocystis* was conjugated by Triparental mating (Elhai et al. 1997). *E. coli* TOP10 cargo cells (carrying pEEK2 plasmids) and *E. coli* HB101 helper cells (carrying pRL443-AmpR conjugative plasmid) were cultivated overnight in LB medium with 50 µg mL^−1^ kanamycin at 37 °C. Cells were collected by centrifugation and resuspended in fresh liquid LB medium without antibiotics. Then, a mixture of wildtype *Synechocystis* (200 μl), cargo cells (1 ml) and helper cells (1 ml), was incubated under 30 µmol photons m^−2^ s^−1^ at 30 °C for 2 hours. The mixture was then spread on a filter (GN-6 Metricel, Pall Lab) placed on a BG11 agar plate without antibiotics for 24 hours of incubation at 30 °C. Individual colonies (Table 1) were selected by transferring the filter onto BG11 agar plates with 50 µg mL^−1^ kanamycin and assessed by PCR using gene-specific primers (Supplementary Table 1).

### DNPH assay for substrate consumption

An assay for determining KICD activity, based on a previously described method Gong et al. (2021), was developed. for KIC and HPP consumption. For the assay calibration, HPP and KIC concentration standards in the range from 0.1 to 3 mM were prepared in the reaction buffer (10 mM MES, 150 mM NaCl, pH 6.0) with 0.5 mM FeSO_4_, 0.5 mM sodium ascorbate, and 1 mM dithiothreitol and measured as triplicates. A blank was prepared from the same buffer that did not contain any substrate. The DNPH assay was started by adding 25 µL of a calibration standard or library screening sample to 25 µL DNPH ethanol solution to a well in a microplate. By contrast, the samples from the activity measurements were mixed in a ratio of 1:1 with the DNPH reagent in a reaction tube. Thereafter, 50 µL of the mixture was added to a well in a microplate. The 96-well plates were then incubated for 40 min at 30 °C which allowed the derivatization of KIC and HPP with DNPH. Next, 200 µL 1M NaOH ethanol solution were added to each well. After another 15 min of incubation at 30 °C, the absorbance at 540 nm was measured in a plate reader (Hidex).

### Assessment of substrate consumption

In vitro activity of all purified KICD variants was measured by determining the KIC and HPP depletion over time using an endpoint assay. Each in vitro reaction contained 4.2 µM enzyme, 2.5 mM of either HPP or KIC in the reaction buffer (10 mM MES, 150 mM NaCl, pH 6.0) with 0.5 mM FeSO_4_, 0.5 mM sodium ascorbate, and 1 mM dithiothreitol. Prior to the start of the reaction the enzymes were incubated at 30 °C for 10 min in the reaction buffer. For the whole experiment, the enzymatic reaction was incubated in a heat block at 30 °C. Samples of the reaction were stopped by mixing them with the acidic 95% ethanol DNPH solution resulting in enzyme denaturation. Triplicate enzymatic reactions were run in parallel and sampled at different time points between 0 and 90 min. Additionally, a negative control (NC) reaction without enzyme and a blank without substrate were prepared and sampled after 0, 30, and 60 minutes. The samples from the reaction replicates were pooled in one mixture with DNPH solution for each timepoint so that it contained equal volumes of reaction sample and DNPH solution. After the sample collection was finished, 50 µL of each sample were separately applied to three wells of a microplate. These technical replicates of the DNPH assay measurements were used to determine an average values standard deviation.

### Cultivation and screening of site specific saturation-mutagenesis libraries

After transformation with one of the mutant plasmid libraries, *E. coli* BL21 (DE3) colonies were picked and used for inoculation of individual pre-cultures in 96-well plates. From each library 73 colonies were selected for a >99% library coverage. Additionally, colonies with the empty vector (pETBB) and the expression vector for wildtype *Rn*KICD were cultivated as negative control (NC) and positive control (PC), respectively. The pre-cultures were cultivated in a polyethylene-sealed 96-well plate with micro-perforations (two holes per well) for gas exchange. Each well contained 200 µL TB and 50 µg mL^-1^ kanamycin. After an overnight incubation at 37 °C and 190 rpm the OD_600_ was measured and averaged for all wells. Each pre-culture was used for the inoculation of one well in a 96-deep-well plate containing 1.3 mL TB medium with 50 µg mL^-1^ kanamycin. The pre-cultures were then supplemented with 40 µL of 85 %, sterile glycerol solution and stored at –80 °C. The main cultures were inoculated to a starting OD_595_ of 0.1 and subsequently cultivated at 37 °C and 190 rpm for 2.5h until an OD_600_ of 0.6-0.8 was reached. Protein production was induced by adding IPTG (final concentration 0.5 mM) to each well, and the cultures were incubated at 25 °C and 250 rpm. for 22 h. Cell growth was determined by measuring at OD_595_. The cells were harvested by centrifuging the plates for 20 min at 2464 x g, 4 °C (Eppendorf, Centrifuge 5810 R). Next, the supernatant was discarded, and the plate dried for 10 min. The plate with the cell pellets was sealed with an adhesive aluminum sheet and stored at –80 °C until used for the screening experiment.

For the lysis the cells in the 96-deep-well plate were thawed on ice. Then 100 µL lysis buffer (10 mM MES, 150 mM NaCl, pH 6.0, B-PER II (Thermo Fisher), 50 U mL^-1^ Benzonase, 1 mg mL^-1^ lysozyme and 1 mM phenylmethylsulphonyl fluoride (PMSF) were added to each well. After incubation for 45 min shaking in RT, the cell debris and insoluble protein were pelleted by centrifuging the 96-deep-well plate for 15 min at 2464 x g and 4 °C. The lysate was kept on ice until the supernatant was used in the subsequent screening assay. For screening of the generated mutant libraries, the DNPH assay was performed as an end-point assay. Thus, the substrate consumption was measured after a predefined time. Before the screening assay, the reaction solutions were prepared with 5 mM of either HPP or KIC of cofactors and substrate. The screening reactions were started for both substrates in sperate 96-well plates by mixing 40 µL cell lysate from each well with 40 µL of the double concentrated reaction solution. In three wells the substrate solution was omitted, and buffer solution without substrate was added instead und used as blank. Hereafter, the two 96-well plates with either HPP or KIC as substrate were incubated at 30 °C for 1h or 3h, respectively. The differential incubation times were chosen due to the lower enzymatic activity of *Rn*KICD with the substrate KIC. After the respective time, 25 µL from each well were transferred to a new 96-well plate in which the DNPH assay was conducted as described above. For the analysis of the results, the substrate consumption in the wells with WT or EVC lysate was averaged. These two average values were used for the comparison of the mutant substrate consumption. Potential hits were selected based on two individual criteria: (1) high KIC consumption and (2) altered consumption for one or both substrates. The first selection criterion was chosen to identify mutants with high KIC conversion rates while the second selection criterion was chosen to investigate the effect on the substrate selectivity. The plasmids of the selected hits were then isolated with the GeneJET plasmid mini prep Kit (Thermo Fisher) for Sanger sequencing (Mix2Seq, Eurofins) with a suitable sequencing primer.

### In vitro isobutene formation

For in vitro isobutene formation, the reaction mixture (2 mL) consisted of 3 mM KIC substrate solution, 1.2% ethanol (as 250 mM KIC stock solution was prepared in absolute ethanol), 0.5 mM sodium ascorbate, 1 mM dithiothreitol, 0.5 mM FeSO_4_, 150 mM NaCl in 10 mM MES buffer pH 6.0. The reaction was initiated by adding purified *Rn*KICD wild-type or mutant enzyme (0.1 mg mL^-1^) to the reaction mixture into 20 mL GC-MS headspace vials (5188-2753, Agilent) sealed with PTFE-lined screw caps (5188-2759, Agilent). The GC-MS vials were incubated at 30 °C for 45 min before isobutene quantification.

### In vivo isobutene formation

For in vivo isobutene production, pre-cultures of *Synechocystis* at OD_750nm_ 0.6 grown in Erlenmeyer flasks were diluted to OD_750nm_ 0.1, then grown to reach OD_750nm_ 0.5-0.6. Then the culture was diluted to OD_750nm_ 0.2 and transferred into 20 mL GC-MS headspace vials (5188-2753, Agilent) filled with 10 mL BG11 medium containing kanamycin (50 μg mL^-1^), and 50 mM NaHCO_3_ (Sigma-Aldrich) was added. The vials were sealed with PTFE-lined screw caps (5188-2759, Agilent), and were shaken horizontally at 120 rpm, under 30 μmol photons m^−2^ s^−1^ at 30 °C.

### Quantification of isobutene in headspace

Samples from in vitro and in vivo experiments were analyzed by a gas-chromatograph (8890, Agilent), equipped with an HP-1ms Ultra I column (19091S-733UI, Agilent) and an inert mass selective detector (5977B, Agilent). The GC-MS method for the isobutene detection was adopted from Rossoni et al. (2015). Prior to injection, the GC vial was incubated for 5 min at 40 °C to equilibrate isobutene concentration in the vial headspace. For detection and quantification, the headspace (100 μL) was injected at an inlet temperature of 150°C and at a split ratio of 100:1. The oven temperature was set to a constant temperature of 40 °C for 3 min followed by a 2 min temperature increase with 30 °C/min. With this method, the retention time was 1.95 min when an authentic isobutene standard (Air Liquide Gas AB) was analyzed. The MS detector was used in selected ion monitoring (SIM) mode to specifically detect the isobutene typical signals at 41 m/z and 56 m/z. Data analysis was done using OpenChrom (Wenig & Odermatt 2010).) and by comparison with the mass spectra and retention time of authentic standard. Three replicate injections were performed per sample.

### Protein extraction from Synechocystis, SDS-PAGE and Western blot

*Synechocystis* cells were harvested by centrifugation at 4,500 x g for 10 min at 4 °C. The cell pellet was washed in 2 mL phosphate buffered saline (1xPBS), centrifuged a 2^nd^ time and resuspended in 200 µL PBS containing 1 mM PMSF (Merck). Acid-washed glass beads (Sigma-Aldrich) were added, and the cells driputed by using a cell homogenizer (Precellys 24, Bertin Instruments). Protein concentration was determined by DC protein assay (Bio-Rad), using albumin from bovine serum (Sigma-Aldrich) as a standard.

Total cell extract of *Synechocystis* (15 µg crude proteins) was separated on SDS-PAGE gels (Any kD Mini-PROTEAN TGX, Bio-Rad), and blotted to PVDF membrane using the Trans-Blot Turbo system (Bio-Rad). The membrane was rinsed with distilled water and blocked with 20 mL of PBS-T0.1 (0.1% Tween 20) containing 3% bovine serum albumin (BSA) for one hour at room temperature with moderate shaking. Then, the membrane was washed three times with PBS-T0.1 for 5 min followed by a 10 min incubation in 10 mL of PBS-T0.1 containing biotin blocking buffer (IBA Lifesciences). Then, anti-Strep-tag II antibody (Strep-Tactin HRP, IBA Lifesciences) was added, and the membrane was incubated for one hour, followed by two washing steps with 20 mL PBS-T0.1 and twice with 20 mL PBS. Subsequently, Strep-tagged proteins were detected using the Clarity ECL substrate (Bio-Rad) according to instruction manual, and quantified using Quantity One Software (Bio-Rad).

### Statistical analysis

All the data are presented as mean ± standard deviation of three independent biological replications. The statistical analysis was performed by Student’s *t*-test, using Excel software (Microsoft). Data were considered significantly different at p < 0.05.

## Acknowledgments

This work was supported by FORMAS—A Swedish Research Council for Sustainable Development (project no. 2021-01669), the Swedish Energy Agency (project no. 44728-1) and the European Union Horizon Europe - the Framework Program for Research and Innovation (2021–2027) under the grant agreement number 101070948 (project PhotoSynH2).

## CRediT authorship contribution statement

CS: Methodology, Investigation, Formal analysis, Validation, Visualization, Writing-Review & Editing. AK: Methodology, Investigation, Formal analysis, Validation, Visualization, Writing-Original Draft, Writing-Review & Editing. BS: Investigation, Formal analysis & Validation. GS: Supervision, Funding Acquisition. HL: Supervision, Visualization, Writing-Review & Editing. CB: Supervision, Writing-Review & Editing. KS: Conceptualization, Supervision, Funding Acquisition, Writing-Review & Editing.

