## Supplemental Figures and Tables for "Structure-guided engineering of α-ketoisocaproate dioxygenase increases isobutene production in *Synechocystis* sp. PCC 6803"

**Supplementary information for Structure-guided engineering of α-ketoisocaproate dioxygenase increases isobutene production in *Synechocystis* sp. PCC 6803**

Conrad Schumann†, Amit Kugler†, Bhavik Shah, Gustav Berggren, Henrik Land, Cecilia Blikstad, Karin Stensjö*

Department of Chemistry-Ångström Laboratory, Uppsala University, SE-751 20, Uppsala, Sweden

*Corresponding author:

† These authors contributed equally.

**Supplementary Figure 1**

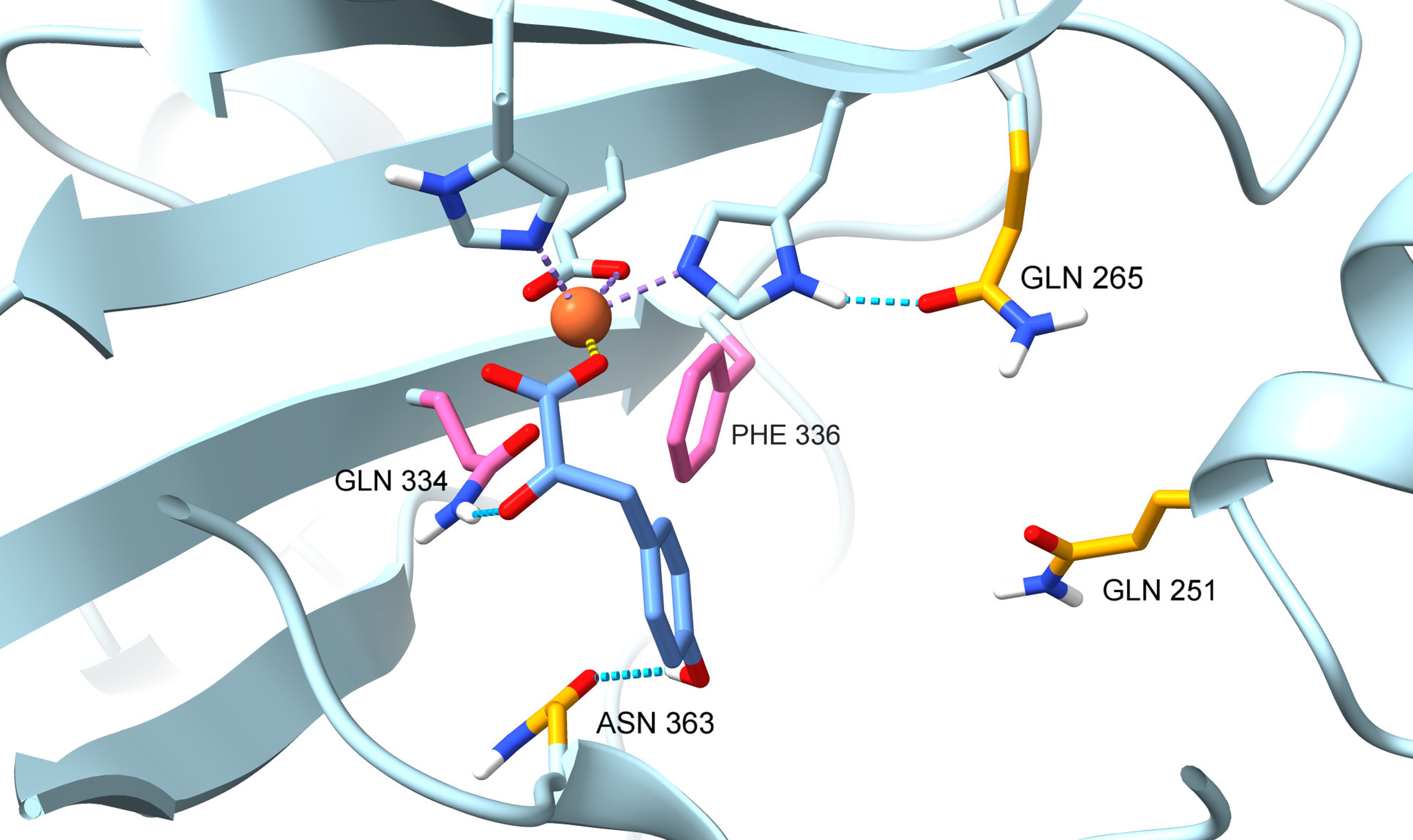

**Supplementary Figure 1** Structural view of the RnKICD homology model active site in open conformation with HPP as substrate. Rational design targets are highlighted in yellow, while site-saturation targets are marked in pink. The structure of HPP is visualized in blue.

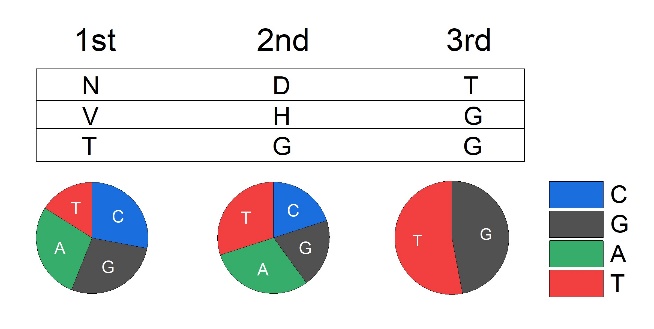

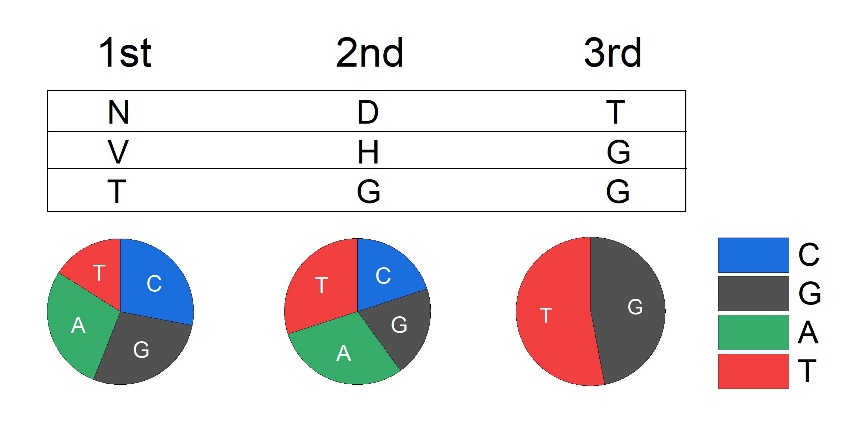
**Supplementary Figure 2**

**A**

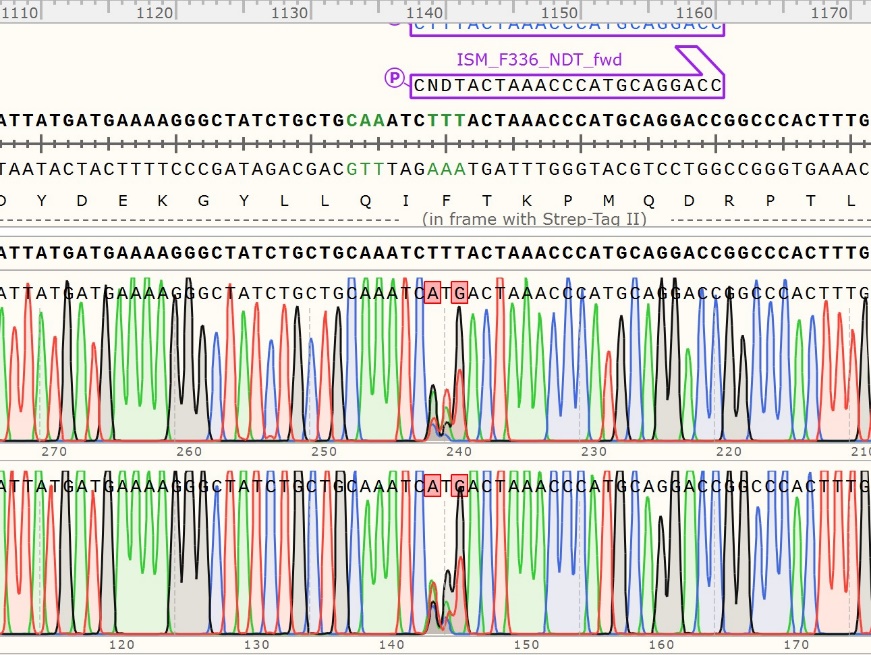

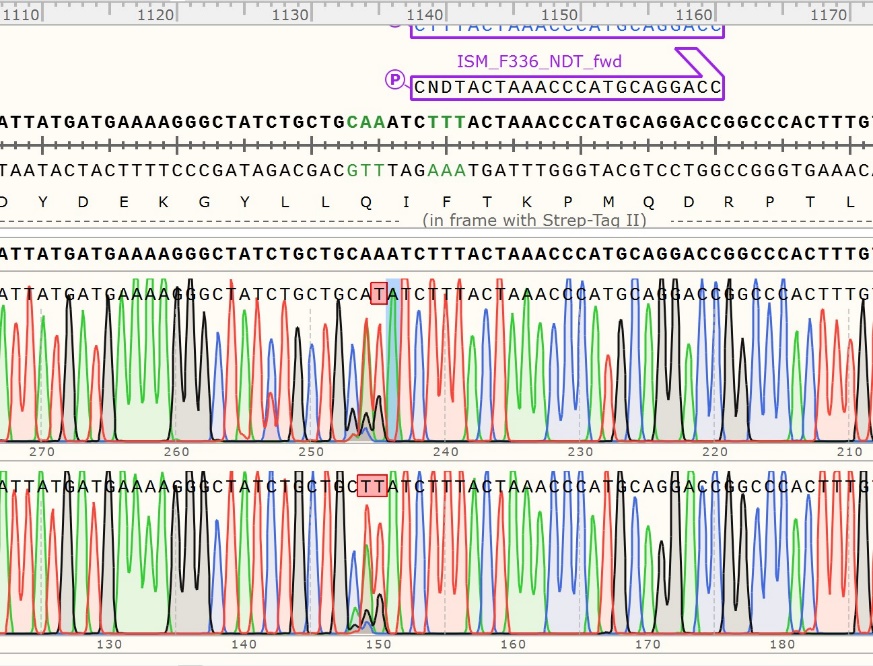

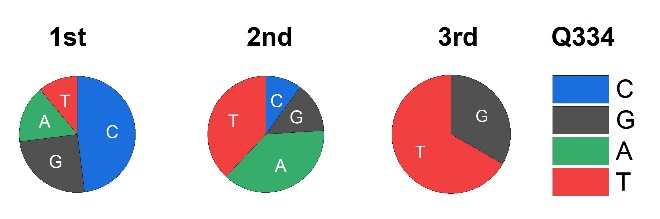

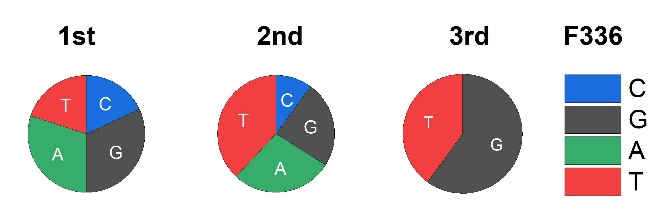

**C**

**B**

**Supplementary Figure 2** Quick quality control (QQC) of libraries generated by site-saturation mutagenesis. A. Degenerate codons and the expected codon distribution according to Kille et al., 2013. B. QQC of site-saturation mutant library on Q334 codon with Sanger sequencing result above and nucleobase distribution below. B. QQC of site-saturation mutant library on F336 codon with Sanger sequencing result above and nucleobase distribution below. The pie charts show the nucleobase distribution in all three codon positions based on estimations from the sequencing results (peak height in the three codon positions). Blue, cytosine/C; black, guanine/G; green, adenine/A; red = thymine/T.

**Supplementary Figure 3**

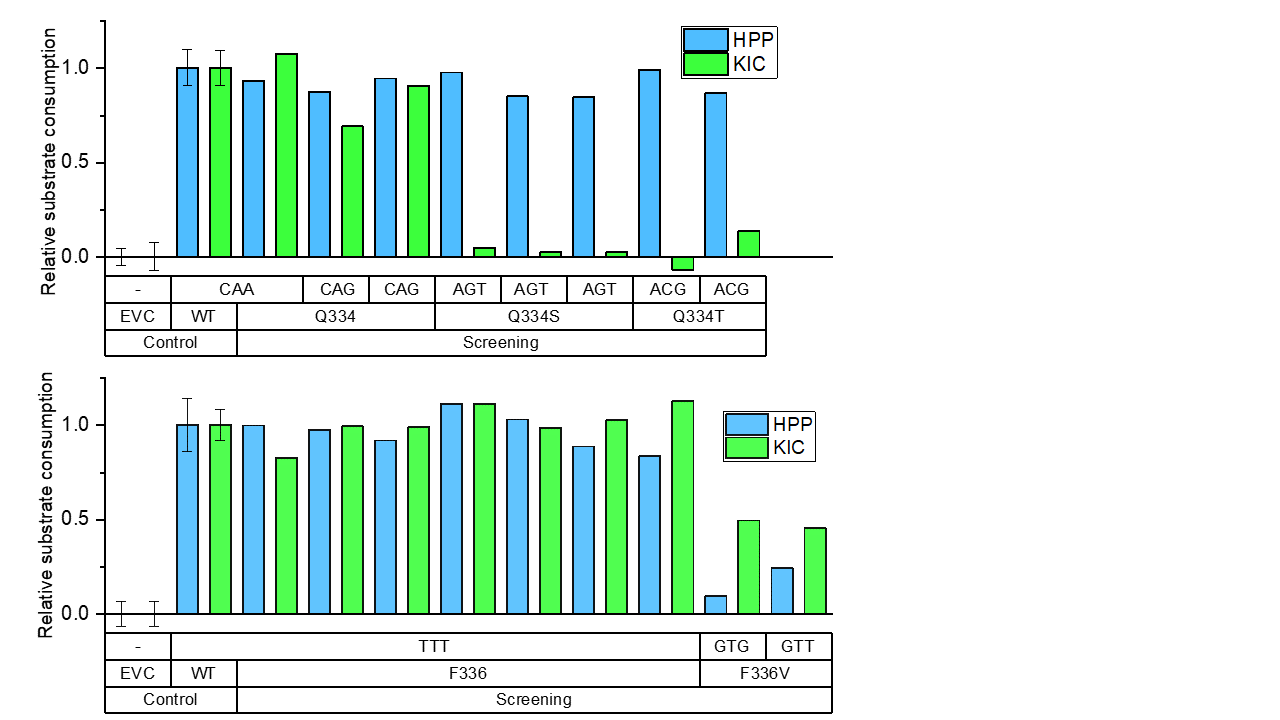

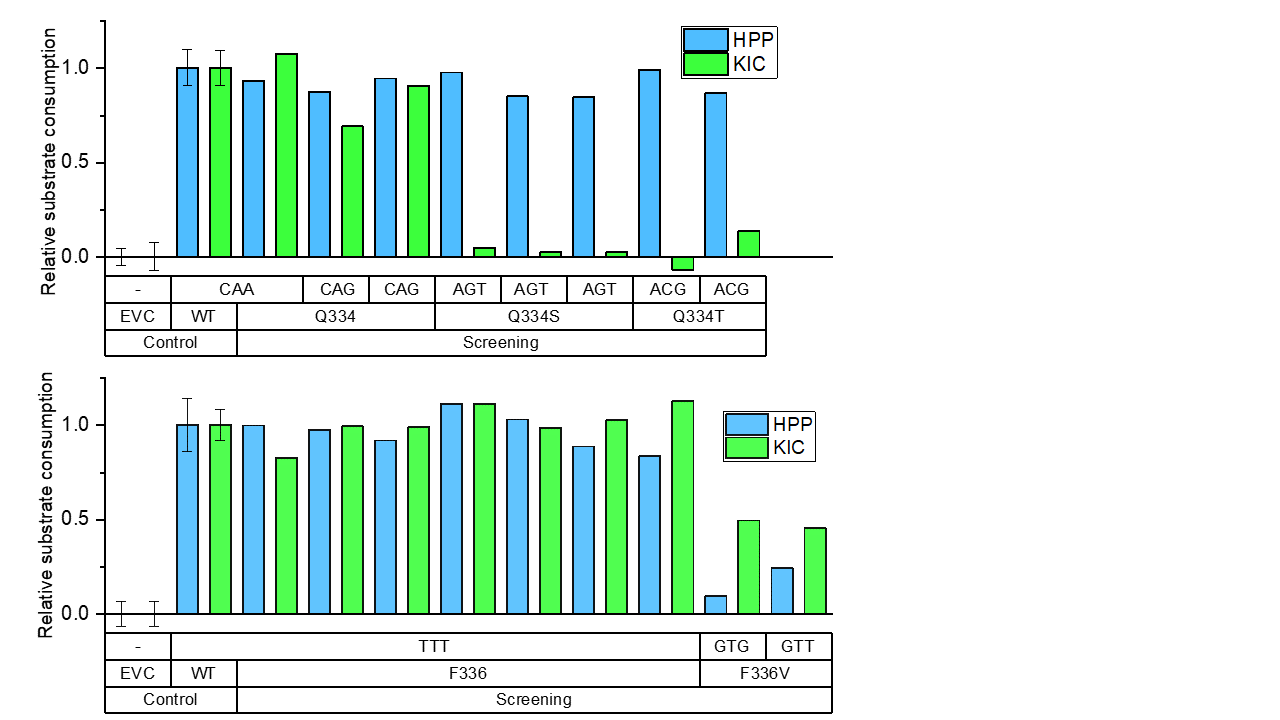

**A**

**B**

**Supplementary Figure 3** Selected hits from the activity screens of transformant libraries generated by site-saturation mutagenesis. A. Screening result for the selected colonies from the Q334X mutant library. Relative consumption rate of KIC (green) and HPP (blue) by Q334X mutants in comparison to WT with the introduced codon and translated amino acid below. B. Screening result for the selected colonies from the F336X mutant library. Relative consumption rate of HPP (blue) and KIC (light green) by F336X mutants in comparison to WT with the introduced codon and translated amino acid below. WT and empty vector control (EVC) for comparison are shown as average with a standard deviation from eight wells per 96-well plate.

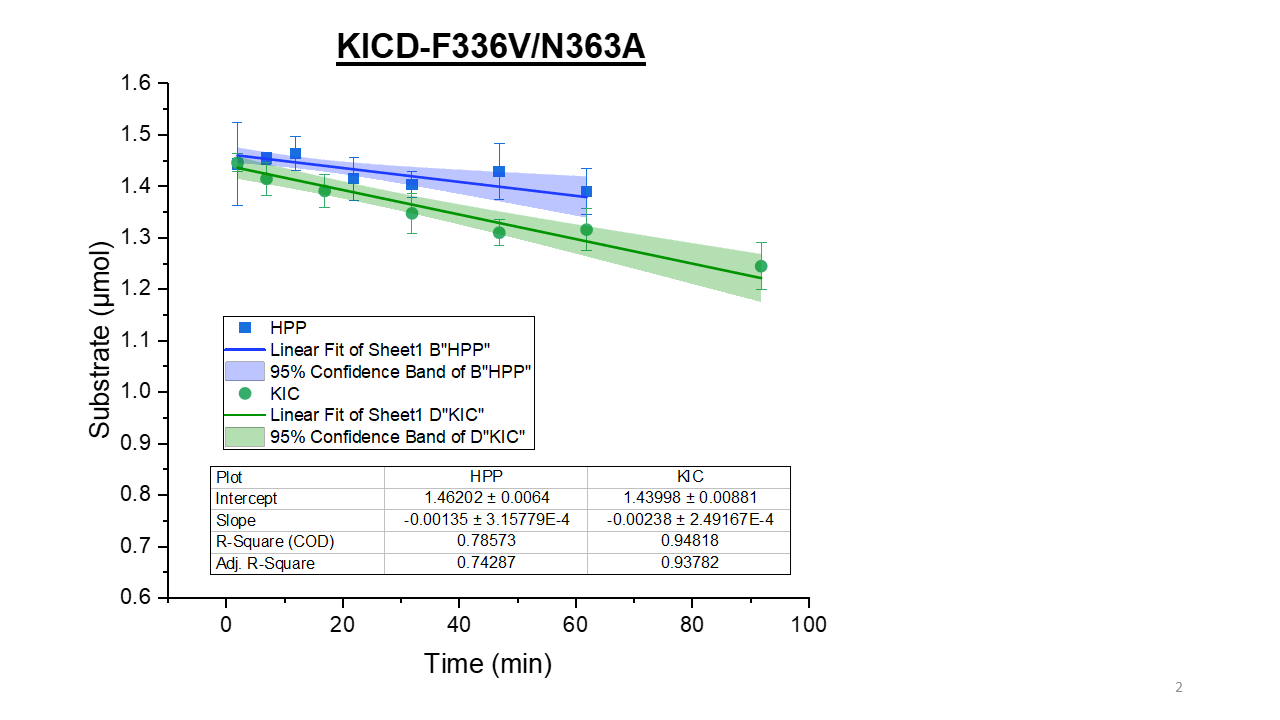

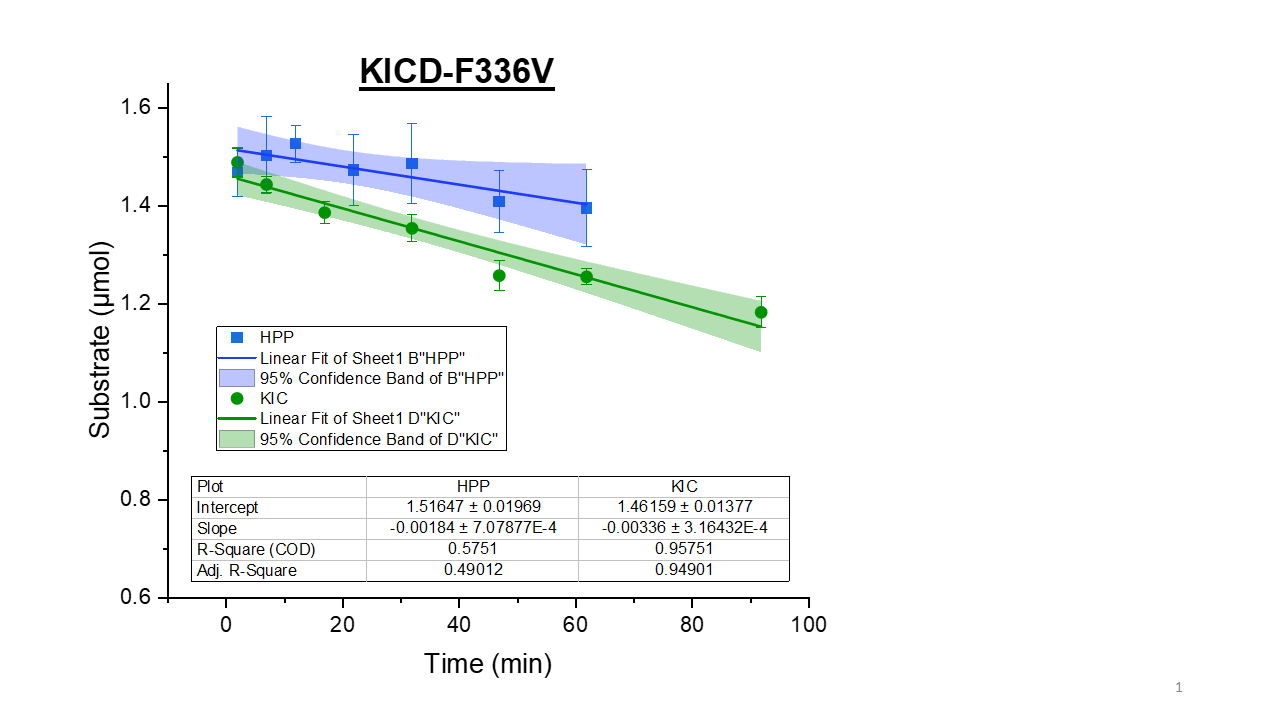

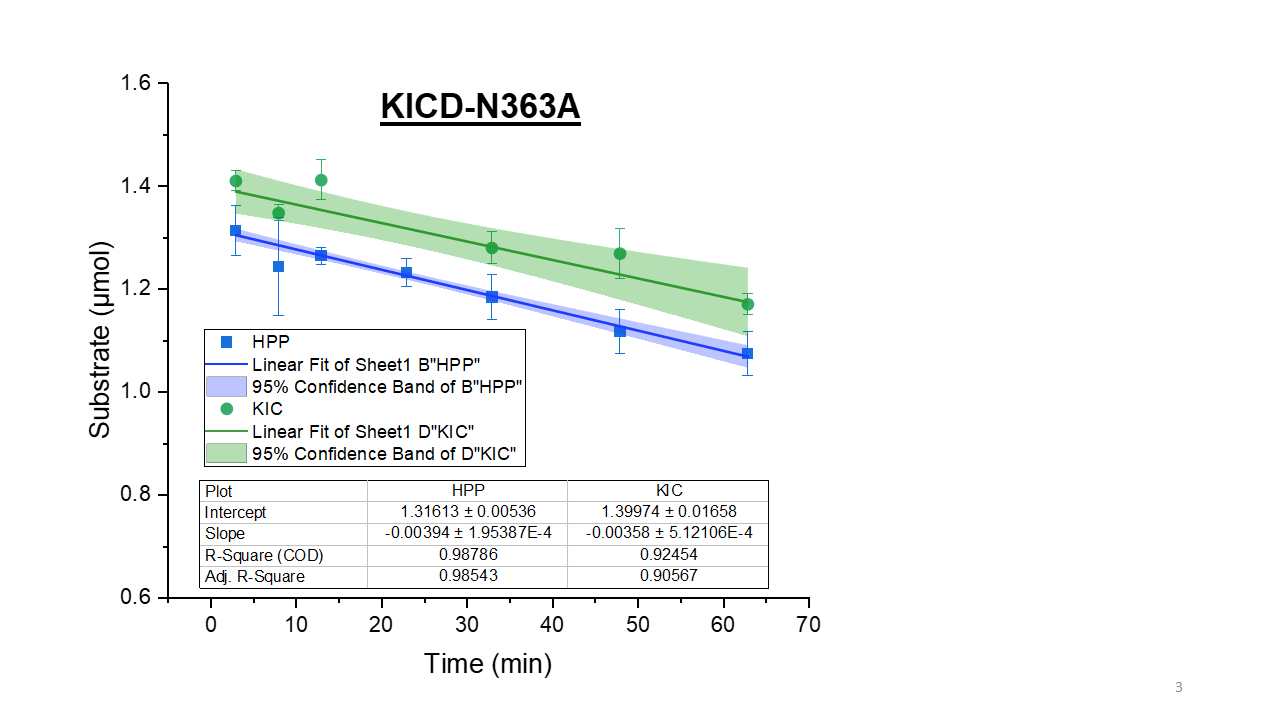

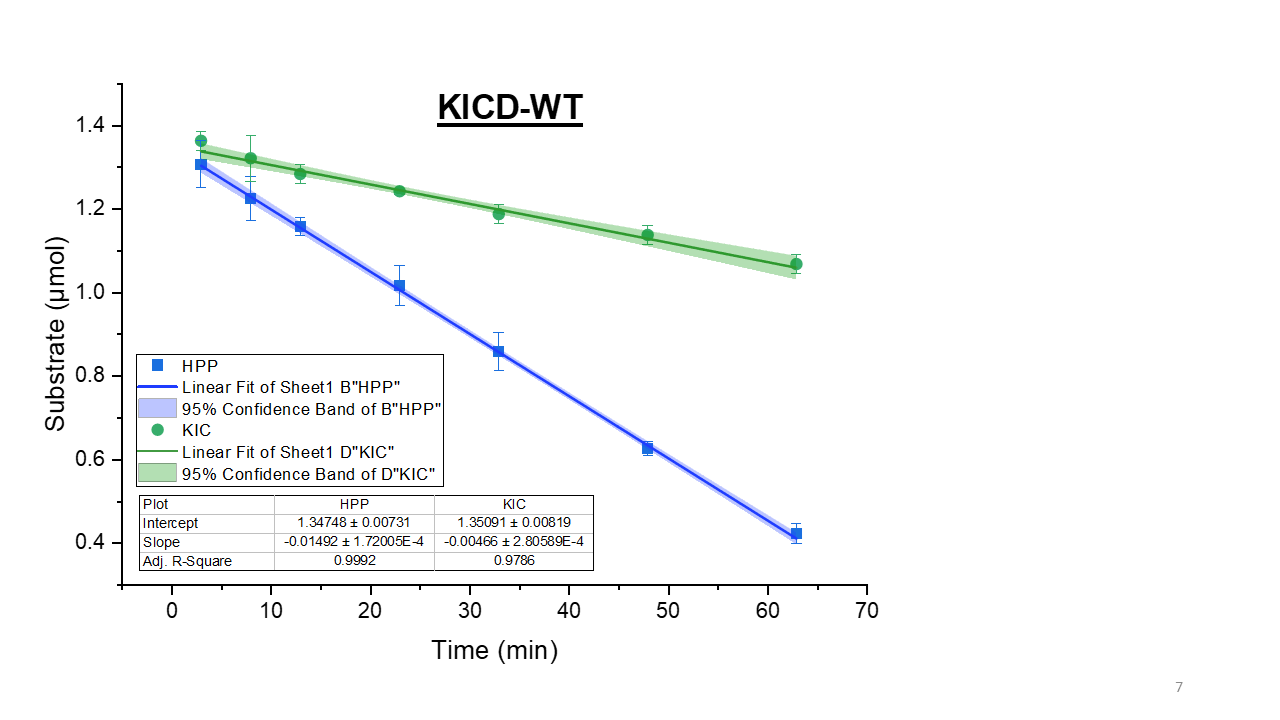
**Supplementary Figure 4**

**F**

**E**

**D**

**C**

**A**

**B**

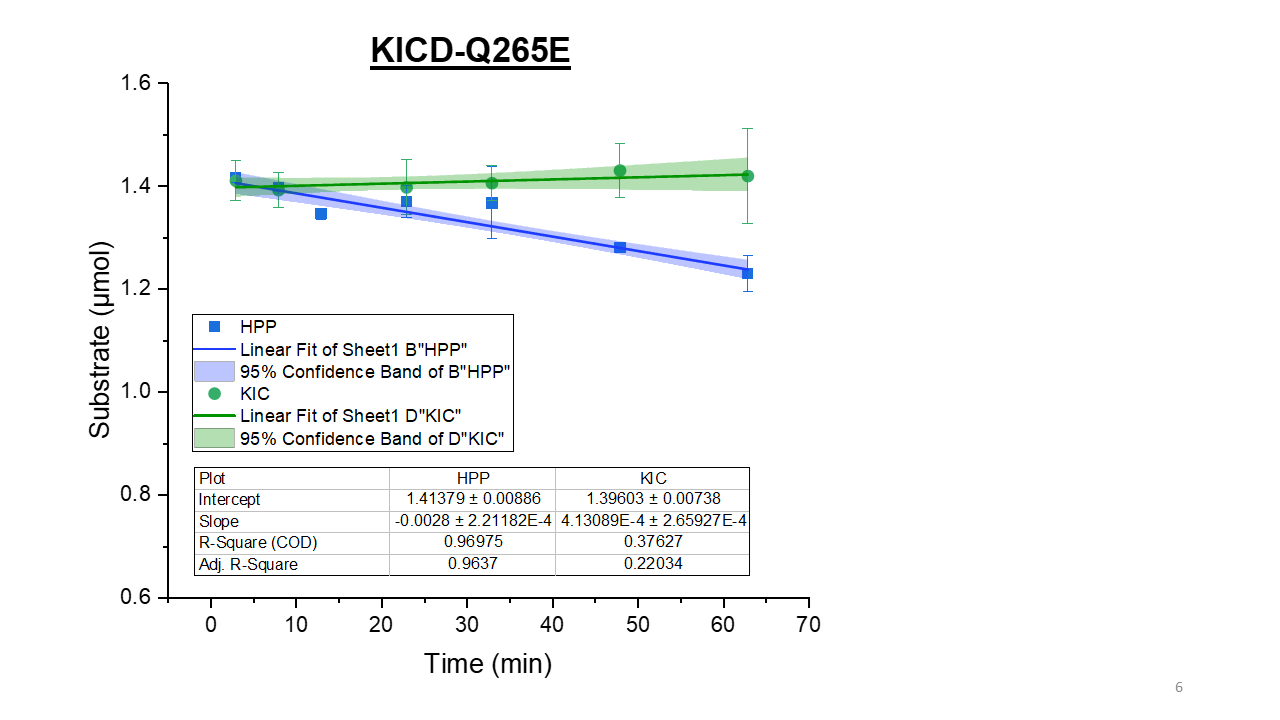

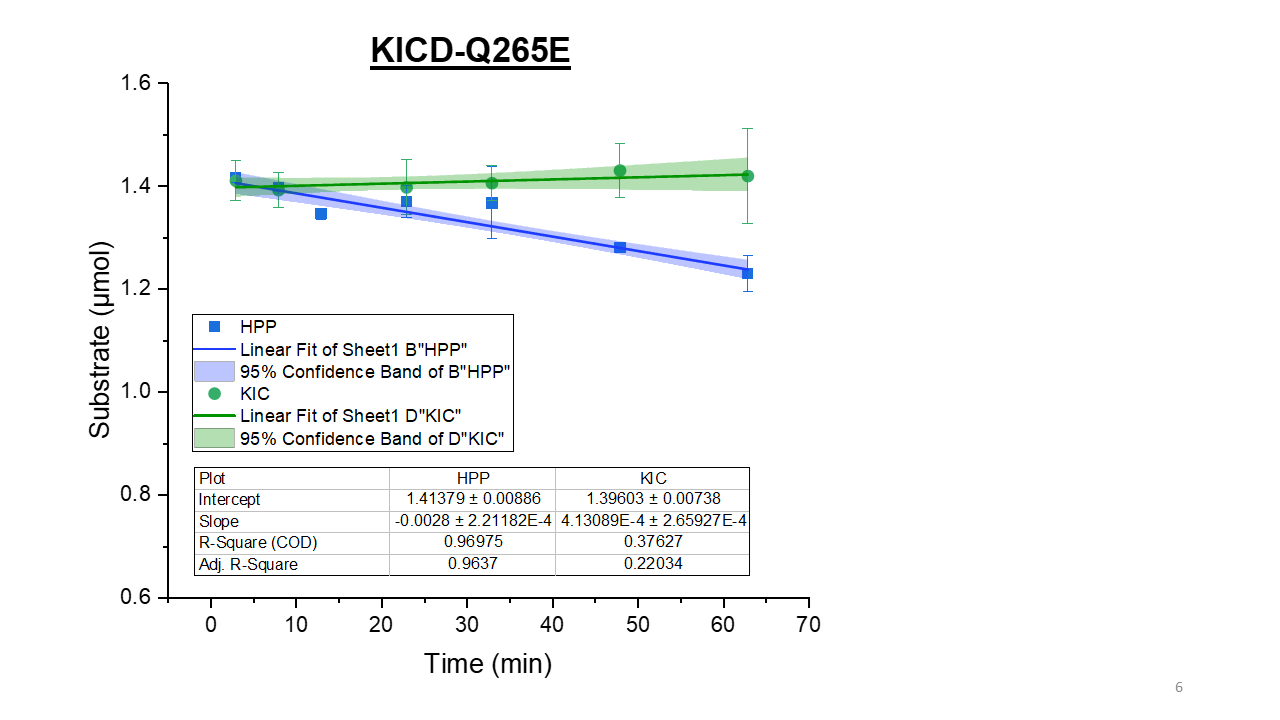

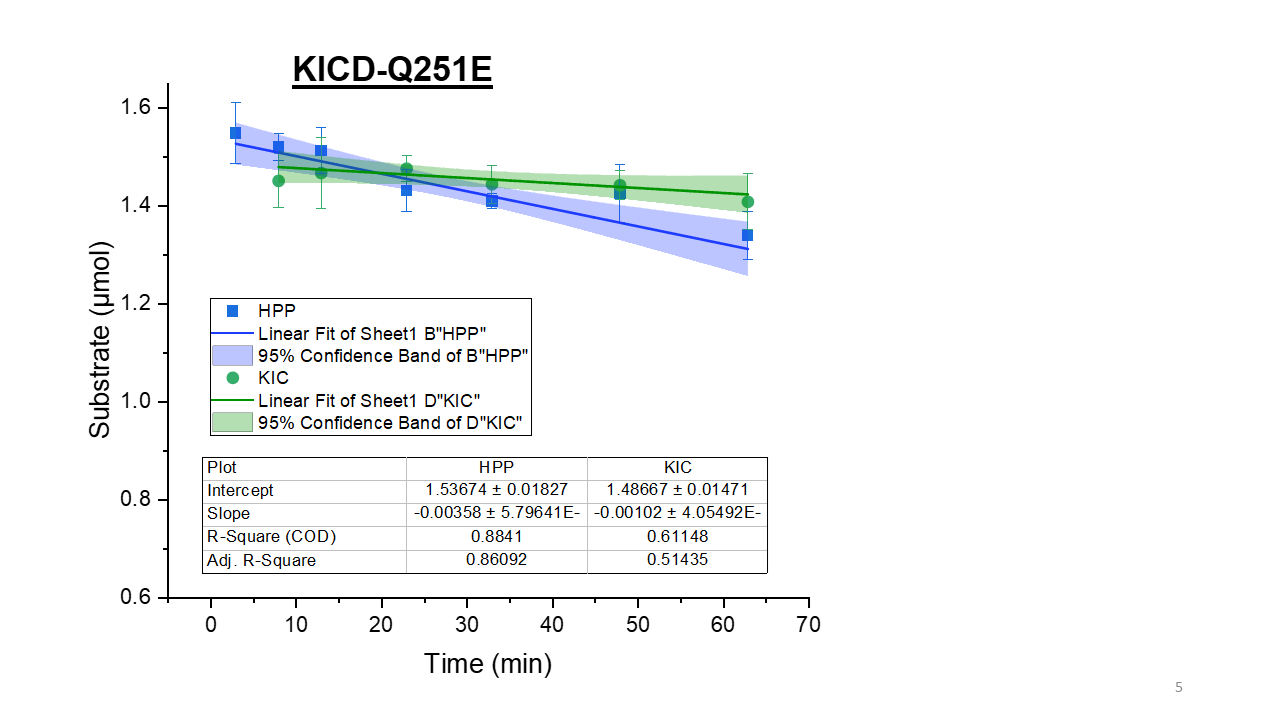

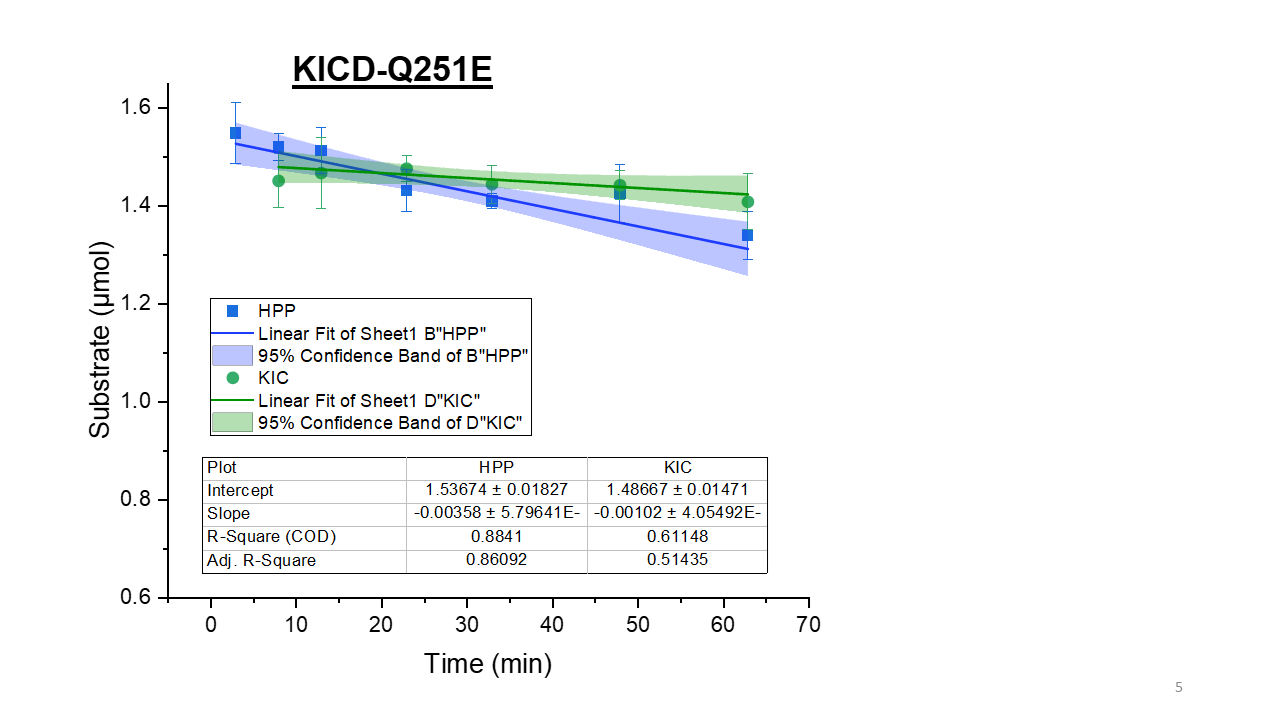

**G**

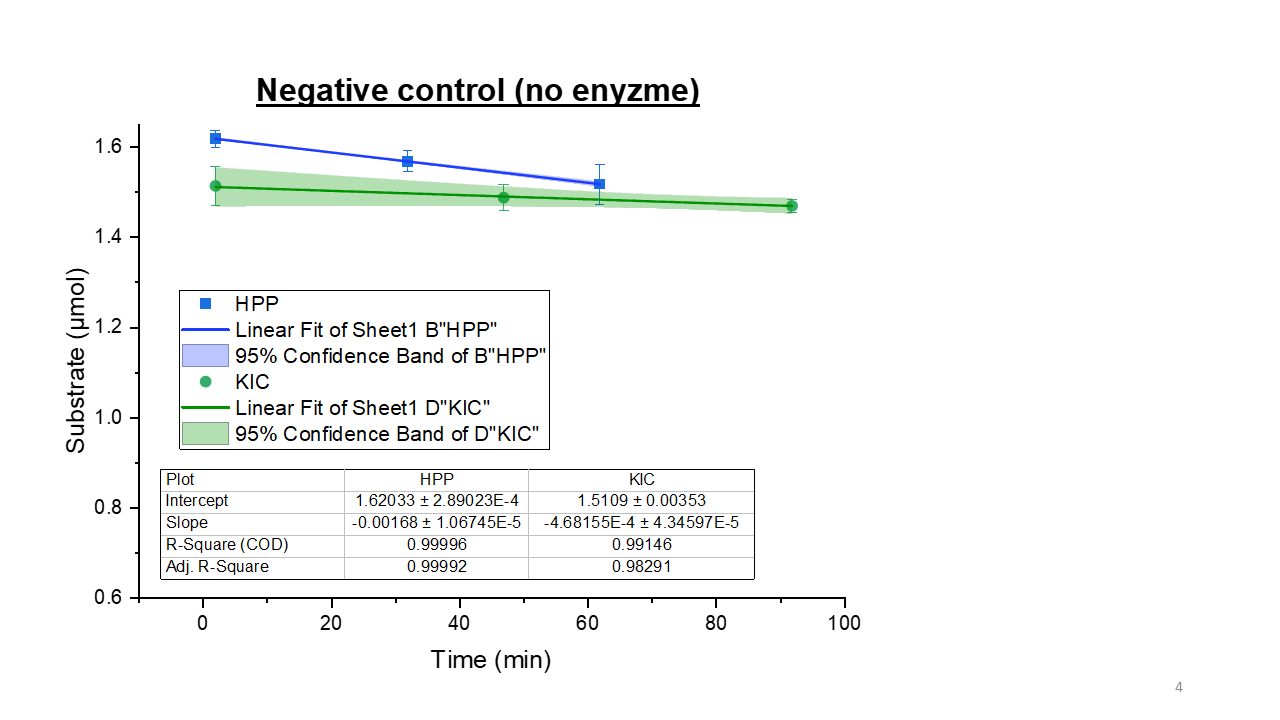

**Supplementary Figure 4**. Progress curves of HPP (blue squares) and KIC (green circles) consumption of purified RnKICD variants at 30°C using DNPH assay. Each point represents mean and standard deviation of three technical replicates containing 4.2 µM enzyme , 2.5 mM substrate (either KIC or HPP), 0.5 mM FeSO_4_, 0.5 mM sodium ascorbate, and 1 mM dithiothreitol in the reaction buffer (10 mM MES, 150 mM NaCl, pH 6.0). The substrate depletion rates and error (in µmol min^-1^) for HPP (blue line and shading) and KIC (green line and shading) are illustrated by linear regressions with 95% confidence bands. A. Substrate depletion of RnKICD-WT. B. Substrate depletion of RnKICD-Q251E. C. Substrate depletion of RnKICD-Q265E. D. Substrate depletion of RnKICD-N363A. E. Substrate depletion of RnKICD-F336V. F. Substrate depletion of RnKICD-F336V/N363A. G. Substrate depletion without enzyme (negative control).

**Supplementary Figure 5**

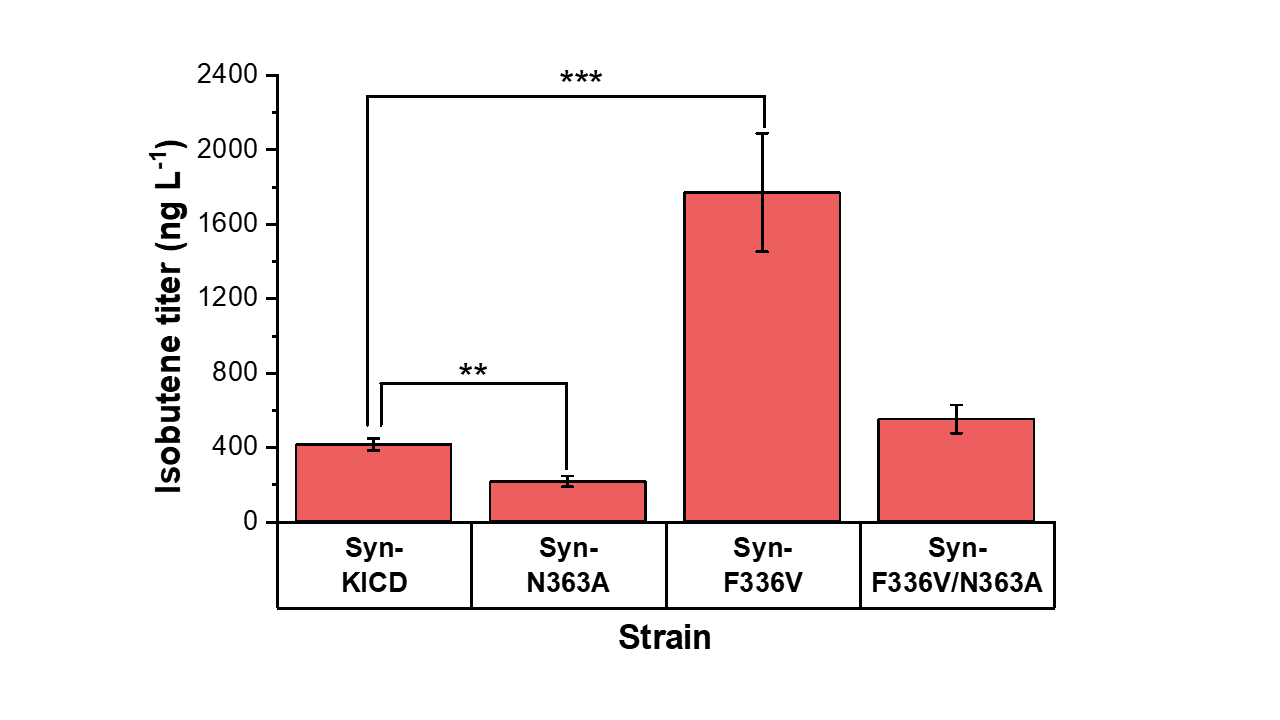

**Supplementary Figure 5**. Isobutene titer (ng mL^-1^) by *Synechocystis* engineered strains after four days of batch cultivation. All the results represent the mean of five biological replicates; error bars represent the standard deviation. Asterisks represent significant differences between the corresponding strain and the base strain, ** p < 0.01, *** p < 0.001 in *t*-test.

**Supplementary Figure 6**

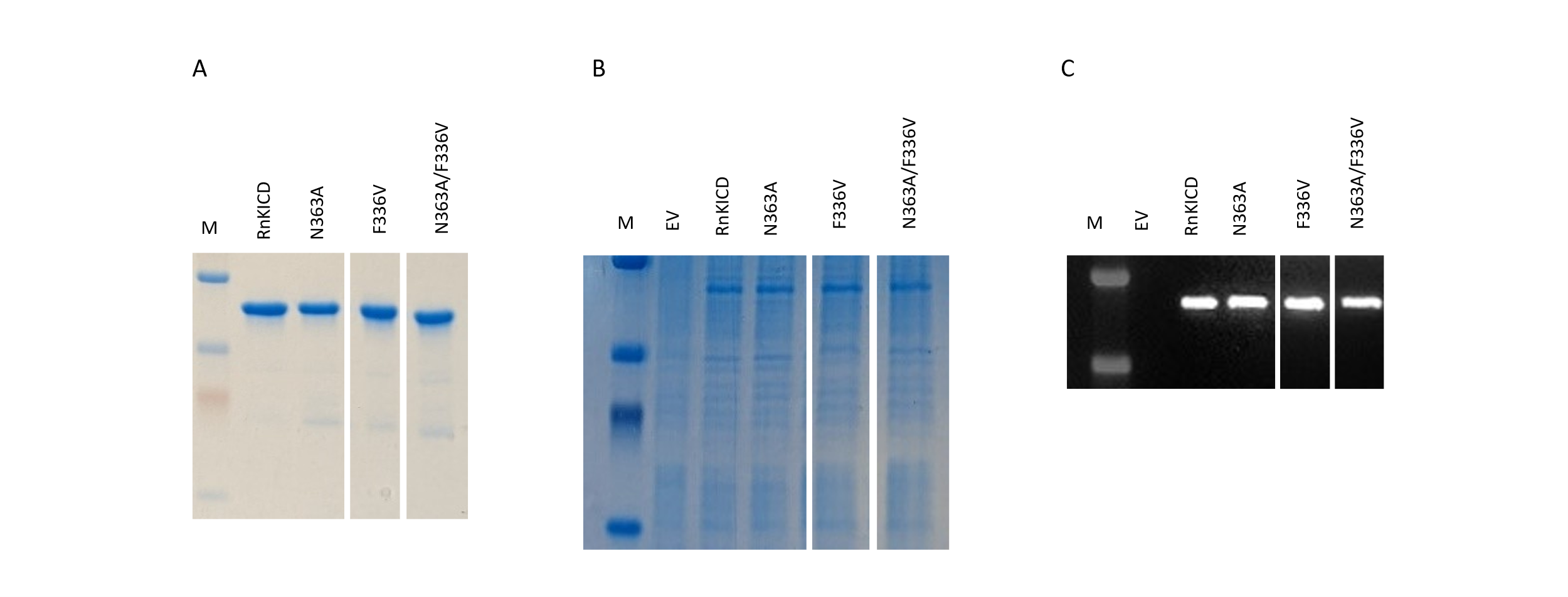

**Supplementary Figure 6** A. SDS-PAGE of purified RnKICD variants. B. SDS-PAGE analysis of recombinant RnKICD variants extracted from *Synechocystis*. C. Western blot analysis of recombinant RnKICD variants extracted from *Synechocystis*. EV is referred to as a *Synechocystis* strain expressing pEEK2 empty vector. M, protein marker.

**Supplementary Table 1** Primers used in this study for site-directed mutagenesis experiment.

| Name | Sequence 5'-3' | Used for |
| --- | --- | --- |
| **pETBB_BB1_FW** | CAAGCGCTCATGAGCCCGAA | General Sequencing |
| **pETBB_BB_RV** | TCGCCAATCCGGATATAGTTCC |  |
| **N363A_FW** | CGCCGGCgctTTTAATTCCTTATTCAAGGCCTTTGAG | N363A mutagenesis |
| **N363A_RV** | GAATTAAAagcGCCGGCGCCAAATCCTTGG |  |
| **Q251E_FW** | GAAAAAATCCgagATTCAAGAGTATGTTGATTATAACGG | Q251E mutagenesis |
| **Q251E_RV** | CTTGAATctcGGATTTTTTCCGGCCAG |  |
| **Q265E_FW** | CGTAgagCATATCGCCCTGCGTACCGAAG | Q265E mutagenesis |
| **Q265E_RV** | GATATGctcTACGCCGGCGCCAC |  |
| **F336V_FW** | GCAAATCgTTACTAAACCCATGCAGG | F336V mutagenesis |
| **F336V_RV** | GTTTAGTAAcGATTTGCAGCAGATAGCCC |  |

**Supplementary Table 2** Primers used in this study for iterative saturation mutagenesis experiment.

| Name | Sequence 5'-3' | Used for |
| --- | --- | --- |
| **ISM_F336_NDT_FW** | /5Phos/CNDTACTAAACCCATGCAGGACC | Generation of F336 site-saturation motagenesis library |
| **ISM_F336_RV** | /5Phos/ATTTGCAGCAGATAGCCCTTTTC |  |
| **ISM_F336_TGG_FW** | /5Phos/CTGGACTAAACCCATGCAGGACC |  |
| **ISM_F336_VHG_FW** | /5Phos/CVHGACTAAACCCATGCAGGACC |  |
| **ISM_Q334_FW** | /5Phos/CTTTACTAAACCCATGCAGGACC | Generation of Q334 site-saturation motagenesis library |
| **ISM_Q334_FW** | /5Phos/ATAHNCAGCAGATAGCCCTTTTCATC |  |
| **ISM_Q334_TGG_RV** | /5Phos/ATCCACAGCAGATAGCCCTTTTC |  |
| **ISM_Q334_VHG_RV** | /5Phos/ATCDBCAGCAGATAGCCCTTTTC |  |
| **ISM_seq_Q334_F336_FW** | ATTACCACTATTCGCCATCTGCGC | Library Sequencing |

**Supplementary Table 3** Comparison of KIC consumption rate at 2.5 mM substrate expressed as specific activity (U mg^-1^) B. Isobutene production rate (ng mg^-1^ min^-1^) at 3 mM KIC also expressed as specific activity (U mg^-1^). U = µmol min^-1^.

|  | **RnKICD (WT)** | **N363A** | **F336V** | **F336V/ N363A** |
| --- | --- | --- | --- | --- |
| KIC consumption rate (U mg^-1^) | 0.035 ± 0.0023 | 0.030 ± 0.0034 | 0.024 ± 0.0029 | 0.016 ± 0.0021 |
| Isobutene production rate (U mg^-1^) | 0.00127 ± 0.00012 | 0.00110 ±  0.00004 | 0.00080 ± 0.00004 | 0.00053 ± 0.00003 |
| Ratio (%) | **3.7 ± 0.4** | **3.7 ± 0.4** | **3.3 ± 0.3** | **3.3 ± 0.4** |

**Supplementary material 1:** Coding sequence of N-terminal-StrepII tagged RnKICD with StrepII highlighted in yellow and glycine-serine linker ([G-S]_3_) highlighted in green.

ATGTGGAGTCATCCTCAGTTCGAGAAGGGTAGCGGAAGTGGATCTATGACTACCTATTCCAACAAGGGACCAAAACCAGAACGGGGGCGTTTTCTCCACTTTCATTCCGTGACTTTTTGGGTTGGTAATGCCAAGCAGGCCGCGAGCTTCTATTGCAACAAAATGGGCTTTGAACCCTTAGCTTATAAAGGCTTGGAAACTGGTAGCCGCGAAGTGGTGAGTCACGTTATCAAACAAGGAAAGATCGTCTTTGTTCTCTGTTCTGCCTTGAATCCCTGGAATAAAGAAATGGGTGATCATCTCGTAAAACACGGAGATGGAGTTAAAGACATTGCCTTCGAAGTGGAAGACTGCGAACATATTGTCCAAAAGGCGCGTGAACGCGGTGCGAAAATTGTTCGAGAGCCATGGGTGGAAGAGGACAAGTTTGGAAAAGTAAAATTTGCCGTGCTTCAAACTTACGGCGATACCACCCATACCCTCGTAGAAAAAATTAATTACACTGGGCGATTTCTGCCGGGCTTTGAAGCCCCCACCTATAAGGATACTTTATTACCCAAGTTGCCATCTTGTAATTTAGAAATTATTGACCATATTGTGGGTAATCAGCCAGATCAGGAAATGGAATCCGCGAGTGAGTGGTACTTAAAAAATTTACAGTTCCATCGGTTTTGGAGTGTGGATGATACCCAGGTGCATACCGAGTACAGTAGTCTCAGGAGCATTGTTGTGGCGAATTATGAAGAATCCATCAAAATGCCGATTAATGAGCCTGCTCCTGGCCGGAAAAAATCCCAAATTCAAGAGTATGTTGATTATAACGGTGGCGCCGGCGTACAGCATATCGCCCTGCGTACCGAAGATATTATTACCACTATTCGCCATCTGCGCGAACGCGGCATGGAATTTTTAGCCGTTCCAAGCAGTTATTATCGTCTCCTACGTGAAAACTTAAAAACTTCCAAGATCCAAGTGAAGGAGAATATGGATGTTCTCGAAGAATTAAAAATCCTTGTGGATTATGATGAAAAGGGCTATCTGCTGCAAATCTTTACTAAACCCATGCAGGACCGGCCCACTTTGTTTTTAGAGGTGATCCAGCGTCATAATCACCAAGGATTTGGCGCCGGCAATTTTAATTCCTTATTCAAGGCCTTTGAGGAAGAACAAGCCTTACGTGGCAATTTGACTGACTTAGAGACCAACGGCGTGAGATCCGGTATGTAA
